# Central pore-opening structure and gating of the mechanosensitive PIEZO1 channel

**DOI:** 10.1101/2023.09.28.559900

**Authors:** Sijia Liu, Xuzhong Yang, Xudong Chen, Xiaochun Zhang, Jinghui Jiang, Li Wang, Jingyi Yuan, Wenhao Liu, Heng Zhou, Kun Wu, Boxue Tian, Xueming Li, Bailong Xiao

## Abstract

PIEZO1 is a mechanically activated cation channel that undergoes force-induced activation and inactivation to control diverse physiological processes^1–4^. However, the distinct functional states and gating transitions remain structurally undefined. In response to changes in membrane forces, PIEZO1 transitions from a curved to a flattened shape, which might lead to opening and subsequent inactivation of the ion-conducting pore^5,6^. Here we employed a PIEZO1 mutant to capture various conformational states including an intermediate state with an opened central pore, allowing us to reveal the gating dynamics of this multiple-gate system. Upon transition from the curved to the partially flattened intermediate state, the trimeric cap undergoes a downward and rotational motion to open the lateral cap-gate, while the loop that connects the cap to the pore-lining inner helix acts as a spring-linker, switching from an extended to a compressed state. Along with the partial flattening of the peripheral blades, these conformational changes collectively dilate the cap-gate, spring-gate and the hydrophobic transmembrane-gate, leading to ion permeation through the transmembrane pore. Upon inactivation, while the blades remain flattened, the cap and spring-linker return to an up and extended state. Molecular dynamics simulations verify the ion-permeating pathway and the distinct ion-conducting states of the assigned close, intermediate, open, and inactivated structures. Mutagenesis and electrophysiological characterizations reveal that domains and residues involved in conformational dynamics of the cap and spring-linker are critical for mechanical activation of PIEZO1. Together, these studies have structurally defined the distinct functional states and the curvature-based activation and inactivation mechanisms of PIEZO1.

## Main

PIEZO1 is a specialized mechanosensor that directly converts membrane tension and curvature into cation permeation^1,6–8^. It is activated within a millisecond (ms) and then followed by fast inactivation^1^, indicating its rapid transition from the closed to open and then inactivated states. Modulation of the gating process might account for the broad spectrum of physiological and pathophysiological roles mediated by PIEZO1^4,9^. For instance, in some cell types^10–12^, the endogenous PIEZO1-mediated currents display drastically slowed inactivation kinetics when compared to heterologously expressed PIEZO1. Furthermore, human disease-causing mutations of PIEZO1 are characterized with slowed inactivation as gain-of-function properties^13,14^. Thus, it is crucial to structurally determine the distinct functional states in order to mechanistically understand the activation and inactivation mechanisms.

Detergent solubilized PIEZO1 forms bowl-shaped trimers comprising a central ion-conducting pore with an extracellular cap and three curved and non-planar blades, each of which contains 38 transmembrane helixes (TMs) and a featured intracellular beam^15–19^. The intrinsically curved structure might represent a closed state as the channel is under a tension-free condition and its hydrophobic TM pore with a radius of 2-4 Å is suggested to be locally unwetted and therefore energetically unfavored for water and ion permeation^20^. By harnessing membrane forces derived from curvature mismatch between the PIEZO1 channel protein and the residing lipid membrane, we have determined the membrane force-induced flattened structure of PIEZO1 reconstituted in liposome vesicles^6^. Flattening the blades leads to a moderate dilation of the TM pore^6^. However, given the presence of constant membrane bending force, the flattened PIEZO1 structure might represent an inactivated state instead of a fully open state. The rapid transition of PIEZO1 from open to inactivated states imposes a daunting challenge to capture the open state structure. Toward this, we have electrophysiologically identified the PIEZO1-S2472E mutant that is stabilized in the conducting state and determined its curved, partially flattened and less resolved but fully flattened structures. Combining structural analyses, molecular dynamics simulations, mutagenesis and electrophysiological characterizations, we have assigned the close, intermediate, open and inactivated states of PIEZO1 and revealed the key structural changes of the dynamic gating process.

### Identification of the S2472E mutant with distinct gating properties

The pore-lining inner helix (IH) of the TM38 of PIEZO1 mainly consists of hydrophobic residues including the pore-facing I2466, L2469 and V2476. The CHAP software prediction and mutagenesis studies suggest that this portion of the TM pore of the curved PIEZO1 structure might form a hydrophobic gate^20–22^. We tested whether mutating the sole polar residue S2472 within this hydrophobic pore segment to negatively charged glutamate (S2472E) might affect channel properties. Using a piezo-driven blunted glass pipette to mechanically probe the cell membrane under a whole-cell recording configuration, we found that S2472E-transfected cells showed higher basal current (Fig. 1a, b), slowed inactivation kinetics with a constant of 104.8 ± 6.2 ms (18.9 ± 2.2 ms for PIEZO1) (Fig. 1a, c) and about 40% of remaining current even after removal of the mechanical stimulus (Fig. 1a, d). Both the mechanically evoked current and Yoda1 (a PIEZO1 chemical activator)^23^ induced Ca^2+^ response were reduced in the S2472E-transfected cells (Extended Data Fig. 1a-f). Consistent with the higher basal current in S2472E-expressing cells, these cells had elevated basal cytosolic Ca^2+^ levels (Extended Data Fig. 1d, e), and were unhealthy and less viable and more difficult to be patched than PIEZO1-transfected cells (Fig. 1d and Extended Data Fig. 1g). In contrast, although the disease-causing mutant R2482H is characterized by slowed inactivation^13^, R2482H-expressing cells had comparable viability as PIEZO1-expressing cells (Fig. 1e). Collectively, these data suggest that some of the S2472E mutant channels might be stabilized in an open state by slowing its transition to the inactivation state, reaching an equilibrium between different functional states. We therefore set out to determine its structure.

**Figure 1.**
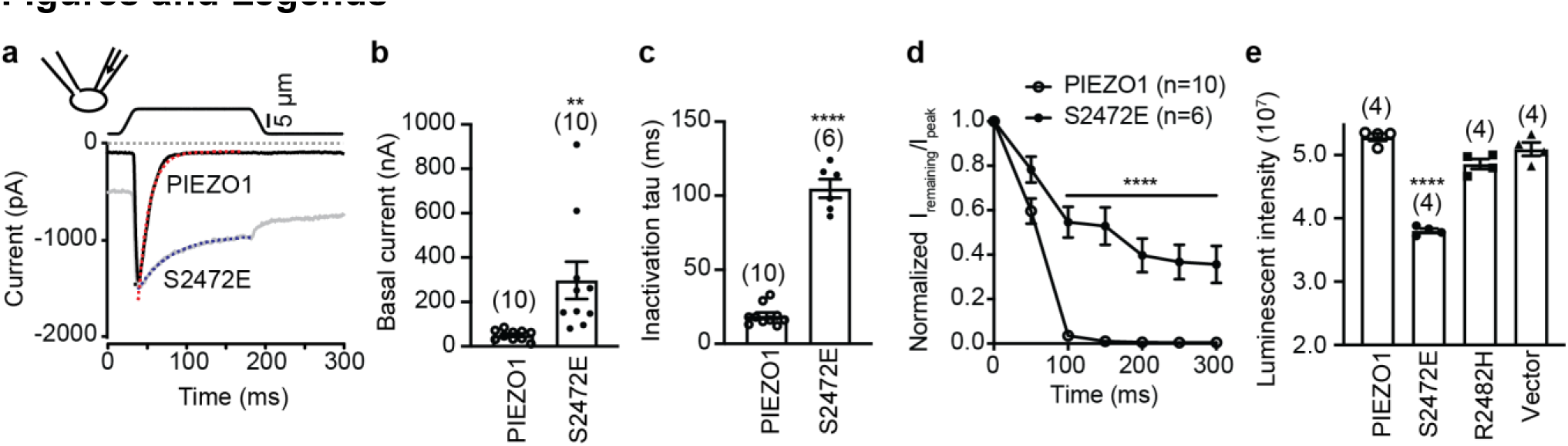
| Functional characterizations of the PIEZO1-S2472E mutant. **a**, Representative poking-evoked whole-cell currents of PIEZO1-KO-HEK293T cells transfected with either wild-type mouse PIEZO1 or the S2472E mutant. The grey dashed line indicates baseline for the current traces, and the red and blue dash lines represent the fitted curve for inactivation Tau calculation. **b**, Scatter plot of the basal current. The recorded cell numbers are labeled. Unpaired Student’s t-test. **P < 0.01. **c**, Scatter plot of the inactivation Tau of the poking-evoked whole-cell currents. The recorded cell numbers are labeled. Unpaired Student’s t-test. ****P < 0.0001. **d**, Normalized ratio of the remaining current at the indicated time points to the peak current. Unpaired Student’s t-test. ****P < 0.0001. **e**, Scatter plot of viable HEK293T cells transfected with the indicated constructs. 4 wells were analyzed for each construct and the experiment was repeated twice. One-way ANOVA with comparison to the PIEZO1 group. ****P < 0.0001.

### Distinct structures of S2472E

As described in detail in the Method and shown in the workflow of single-particle cryo-EM (Extended Data Fig. 2-5), we have overcome the relatively low yield of protein purification of S2472E and determined its various conformational structures. The initial imaging process of the three collected datasets led to a curved structure of S2472E at an overall resolution of 4.07 Å, which largely resembles the curved structures of PIEZO1 (dubbed PIEZO1-Curved) determined in detergent (EMD-6865) (Fig. 2a and Extended Data Fig. 2, 3). We term this structure S2472E-Curved.

**Figure 2.**
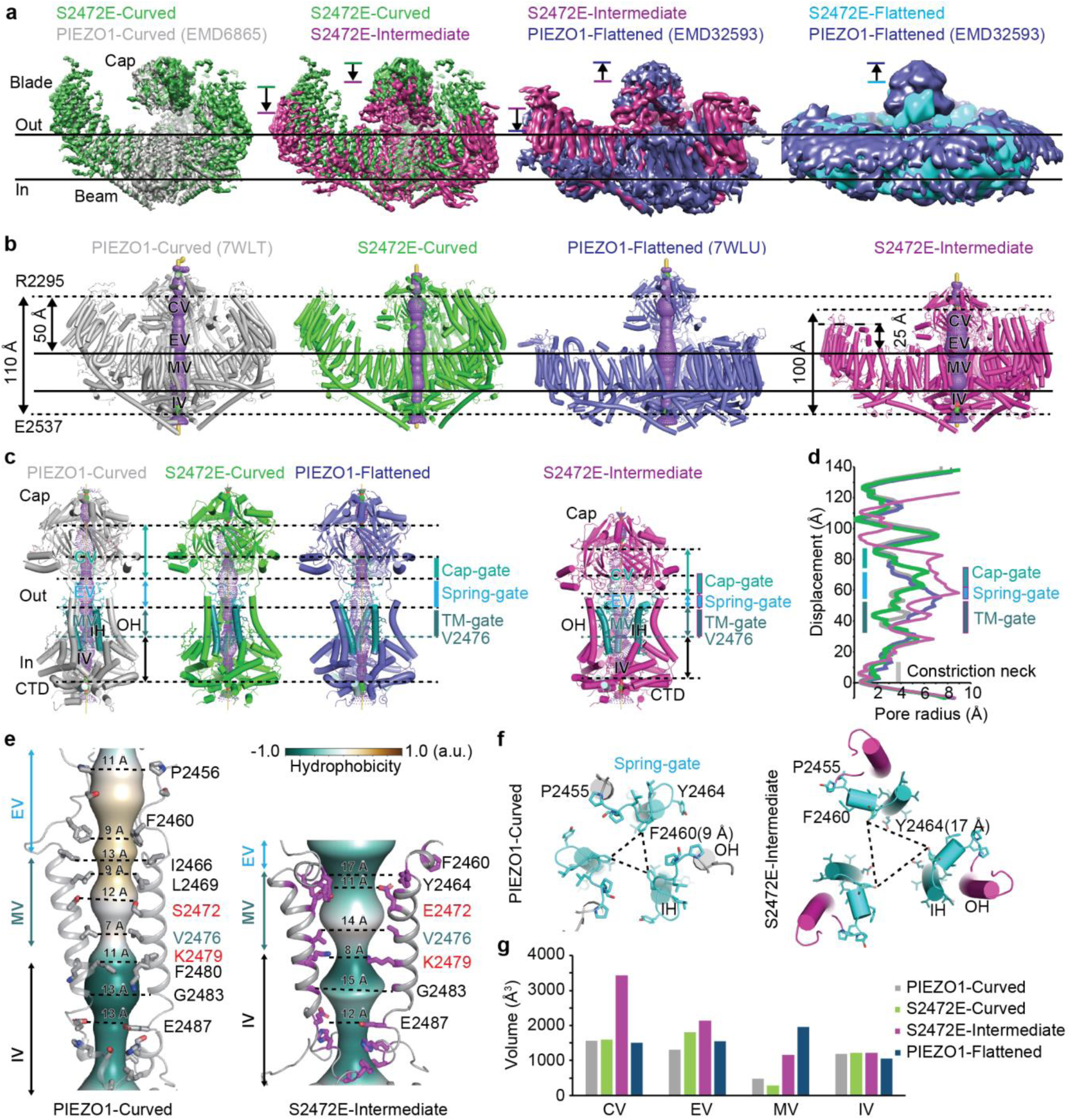
| Distinct structural states. **a**, Comparison of the three distinct structure maps of S2472E (S2472E-Curved, -Intermediate and -Flattened) (at a contour level of 8.19σ, 9.68σ and 6.16σ) with the PIEZO1-Curved (EMD-6865) and PIEZO1-Flattened (EMD-32593) maps. The S2472E-Intermediate map was low-pass filtered to 7 Å in the third column. S2472E-Flattened and PIEZO1-Flattened maps were low-pass filtered to 10 Å in the fourth column. The motion of the cap and blade is indicated. **b**, The side view of the indicated structures. The central pore is generated by the program HOLE and shown in spheres (pore radius: red, < 1 Å; green, < 2 Å; purple, > 2 Å). The top and bottom black lines indicate the outer and inner sides of the planar lipid bilayer. The top and bottom dashed lines indicate the positions of the extracellular constriction site R2295 in the extracellular cap, and the intracellular constriction site E2537 in the intracellular C-terminal domain (CTD), respectively. The CV, EV, MV, IV stand for cap vestibule, extracellular vestibule, membrane vestibule, and inner vestibule respectively. In the S2472E-Intermediate structure, the lowered position of R2295 is labeled using a dashed line. The vertical distance between R2295 and E2537 along the central pore axis and the height of the blade relative to the planar outer membrane are measured and labeled. **c**, The aligned side view of the central pore module of the indicated structures. The central pore is generated by the program HOLE and shown in licorice (pore radius: red, < 1 Å; green, < 2 Å; purple, > 2 Å). T2332/V2333/E2334/Y2335 shown in green cyan constitute the cap-gate region; P2455-G2465 shown in cyan constitute the spring-linker; I2466-F2485 shown in deepteel constitute the inner helix (IH) and the transmembrane (TM) gate. **d**, Pore radius along the central axis of the indicated structures: PIEZO1-Curved (gray), S2472E-Curved (green), PIEZO1-Flattened (slate); S2472E-Intermediate (magenta). Regions corresponding to the cap-gate, spring-gate and TM-gate are labeled with green cyan, cyan and deeptool bars, respectively. **e**, Side view of the spring-linker and IH-enclosed EV, MV and IV (2 out of 3 subunits presented for clarity) showing the hydrophobicity profile generated by CHAP and the side chain distance of the PIEZO1-Curved (left) and S2472E-Intermediate (right). **f**, Zoomed top view at the spring-gate of the PIEZO1-Curved (gray) (upper) and S2472E-Intermediate (magenta) (below). The distance between F2460 and that of Y2464 is measured. **g**, Measured volumes of CV, EV, MV and IV from the indicated structures.

We noticed that some 2D classes of the S2472E had flatter blades (Extended Data Fig. 2e), prompting us to carry out template-based auto-picking of those particles with flatter blades and multi-reference 3D classification, resulting in 4 classes of structures showing partially (class 1) and likely fully flattened blades (class 2) (Extended Data Fig. 5). Based on class 1, a partially flattened structure of S2472E was eventually determined at an overall resolution of 4.00 Å, whose blades are notably lower than the curved structures of both PIEZO1 and S2472E, but higher than the fully flattened structure of PIEZO1 (EMD-32593, termed PIEZO1-Flattened) (Fig. 2a and Extended Data Fig. 4, 5). We termed this intermediately flattened structure S2472E-Intermediate.

Despite extensive 3D classification, the resolution of the class 2 map could not be substantially improved (Extended Data Fig. 5). Nevertheless, when overlayed with the map of the PIEZO1-Flattened, the blade region of this map is embedded in the fully flattened blade region of the PIEZO1-Flattened map, while the cap region appears to be lower (Fig. 2a). Thus, we term this fully flattened map the S2472E-Flattened, which might represent a conformational state distinct from both the S2472E-Intermediate and the PIEZO1-Flattened structure.

We built the models for the S2472E-Curved and S2472E-Intermediate structures with bound phospholipids. The S2472E-Curved structure forms a homotrimer with a similar pore radius as the PIEZO1-Curved structure (PDB: 7WLT and 5Z10) (Extended Data Fig. 6a, b), in which the 26 resolved TMs (12 TMs in the N-terminal region unresolved) curved outwardly from the center to the periphery with a height of ∼50 Å above the planar membrane (Fig. 2b). A top-down view shows that the TM blade region of the S2472E-Curved is slightly expanded when compared to the PIEZO1-Curved (Extended Data Fig. 6a, b), which might be induced by the S2472E mutation. Some bound phospholipids were resolved (Extended Data Fig. 2h, i and Extended Data Fig. 4g, h). The overall resemblance of the S2472E-Curved to the PIEZO1-Curved structure suggests that the S2472E mutation did not cause drastic changes in the overall structure.

Compared to the PIEZO1-Curved or S2472E-Curved structures (Fig. 2a, b and Extended Data Fig. 6c and Supplementary Video 1), the structural model of the partially flattened S2472E-Intermediate shows that the peripheral blades are partially flattened for ∼25 Å and moderately expanded, the intracellular beams are bent at the pivot position of S1341, L1342 and L1345 (Fig. 2b and Fig. 3c), and the top cap is lowered toward the planar membrane for ∼10 Å, which leads to a shortening of the vertical distance between the constriction residues R2295 in the extracellular cap and E2537 at the intracellular C-terminal domain along the central axis from 110 Å to 100 Å (Fig. 2b). A top-down view shows that the cap and blades show an anti-clockwise motion from the curved to the partially flattened structure of S2472E (Extended Data Fig. 6c and Supplementary Video 2).

**Figure 3.**
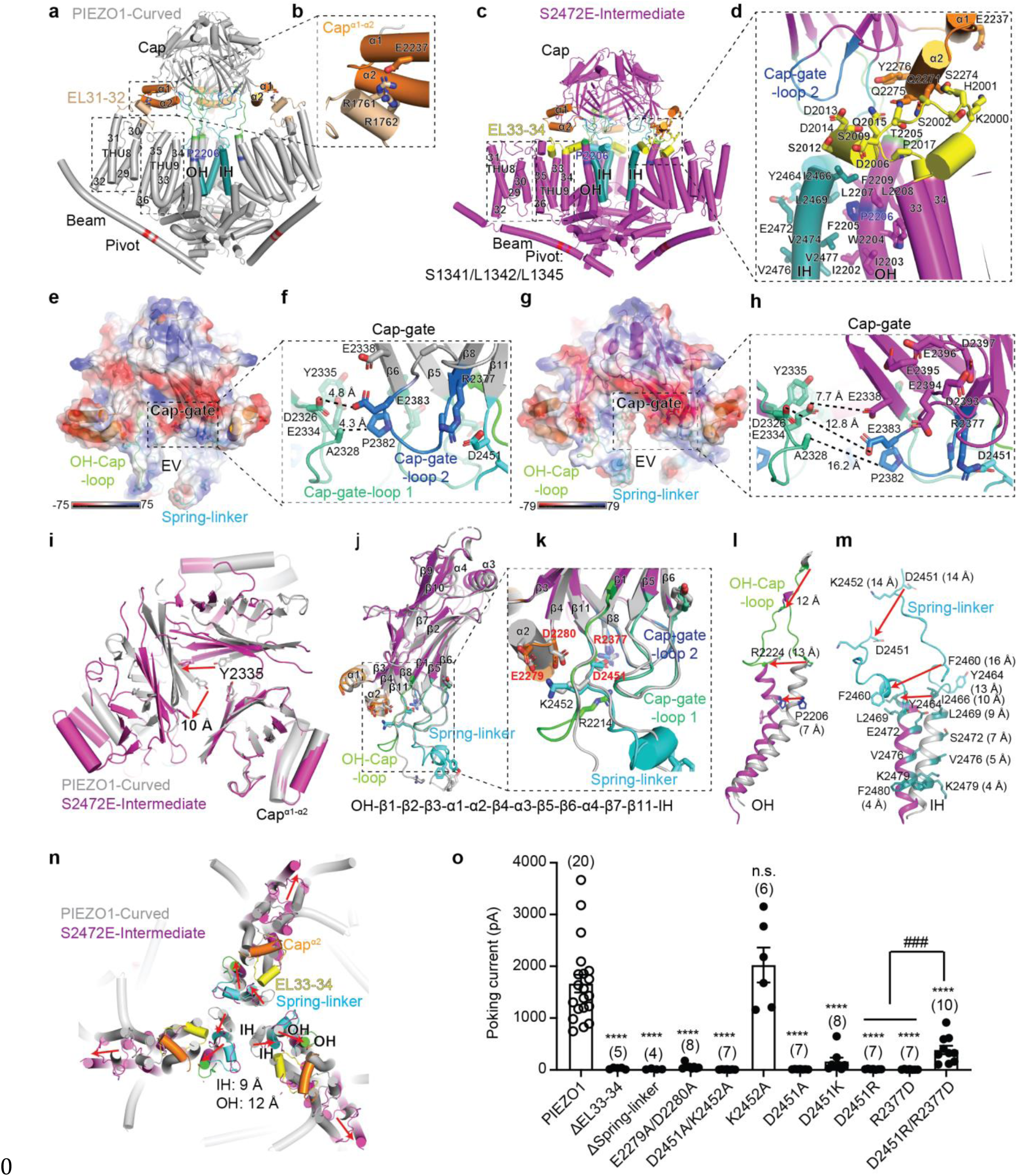
| Gating of the cap-, spring- and TM-gates. **a** and **c**, Side view of the central region of PIEZO1-Curved (gray) (**a**) and S2472E-Intermediate (magenta) (**c**) containing the Beam, THU8 (TM29-32), THU9 (TM33-36), and the pore module. The pivot residues S1341, L1342 and L1345 in the Beam and the kink residue P2206 in the OH are labeled. **b** and **d**, Enlarged view showing the interaction between the Cap^α1^ and the EL31-32 of THU8 in the PIEZO1-Curved (**b**) or between the Cap^α2^ and the EL33-34 of THU9 in the S2472E-Intermediate (**d**). Hydrophobic residues in OH and IH that mediate the extensive hydrophobic interaction are labeled. **e** and **g**, Electrostatic surface potential and model of the cap of the PIEZO1-Curved (**e**) and S2472E-Intermediate structures (**g**). **f** and **h**, Enlarged view of the cap-gate composed of the cap-gate-loop 1 and 2 of the PIEZO1-Curved (**f**) and S2472E-Intermediate structures (**h**). The side chain distance of the labeled residues is labeled. **i**, Superimposed top-down view zoomed at the cap-gate residue Y2335 of the cap of the PIEZO1-Curved (gray) and S2472E-Intermediate (magenta) structures aligned at the Cap^α1-α2^ of one subunit. Red arrows indicate the displacement of the Y2335 residues in the other two subunits. **j**, Superimposed side view of one subunit of the cap domain of the PIEZO1-Curved (grey), S2472E-Intermediate (magenta and the differentially colored domains) structures aligned at the Cap^α1-α2^ of one subunit. **k**, Enlarged view of the upper portion of the spring-linker with the residues involved in electrostatic interactions labeled. **l**, Superimposed side view of OH and OH-Cap-loop of the PIEZO1-Curved (gray) and S2472E-Intermediate (magenta) structures. The displacement is indicated by the red arrows. **m**, Superimposed side view of the spring-linker and IH of the PIEZO1-Curved (gray) and S2472E-Intermediate (magenta) structures. The displacement is indicated by the red arrows. **n**, Superimposed top-down view of the TM regions of the PIEZO1-Curved (gray) and S2472E -Intermediate (magenta and the differentially colored domains) structures aligned at their CTD. Red arrows indicate the displacement. **o**, Scatter plot of the poking-evoked whole-cell currents from PIEZO1-KO-HEK293T cells transfected with the indicated constructs. The recorded cell numbers are labeled. *indicates One-way ANOVA with comparison to the PIEZO1 group. ****P < 0.0001. ^#^ indicates One-way ANOVA comparison of D2451R or R2377D with D2451R/R2377D. ^###^P < 0.001.

Relative to the S2472E-Intermediate structure, the blades of the PIEZO1-Flattened are further flattened for ∼25 Å, but its cap has an upward and clock-wise motion back to the upstate that is similarly observed in the curved structures (Fig. 2a, b). On the basis of the method shown in Extended Data Fig. 7, the measured middle-plane curvature radii of the PIEZO1-Curved (PDB: 7WLT, 5Z10 and 6B3R), S2472E-Curved, S2472E-Intermediate and PIEZO1-Flattened structures are ∼10-12 nm, 14 nm, 32 nm, 117 nm, respectively (Fig. 6). These analyses demonstrate that S2472E-Intermediate represents a conformational state distinct from the previously determined PIEZO1-Curved and PIEZO1-Flattened structures.

### Dilation of the central ion-conducting pore

HOLE-based pore analysis of the PIEZO1 and S2472E structures shows a continuous central pore along the extracellular Cap, the IH-enclosed pore, and the intracellular C-terminal domain (Fig. 2b). Compared to the PIEZO1-Curved and S2472E-Curved structures, the outer portion of the IH of the S2472E-Intermediate structure is laterally expanded, increasing the hydrophobic TM pore radius (Fig. 2c, d). For instance, the diagonal distances between the side chains of the hydrophobic gate residue V2476 are increased from 7 Å to 14 Å (Fig. 2e), exceeding the size barrier of ∼9-12 Å required for wetting and ion permeation of hydrophobic pore^24^. Additionally, the hydrophobic pore region is flanked with hydrophilic residues Y2464 and E2472 at the top side and K2479 at the bottom (Fig. 2e). CHAP-based hydrophobicity analysis shows that the TM pore of the S2472E-Intermediate is largely hydrophilic, in sharp contrast to the hydrophobic pore of the PIEZO1-Curved structure (Fig. 2e).

Compared to the PIEZO1-Curved and S2472E-Curved structures, the S2472E-Intermediate structure shows an apparent enlargement of the bottom portion of the cap vestibule (CV) and the extracellular vestibule (EV) that is enclosed by the three linkers connecting the cap to the three IHs (Fig. 2b-e). As described in more detail below, we term these two portions as cap-gate and spring-gate, respectively, whose radii are enlarged from ∼2-5 Å to 6-9 Å upon transition from the PIEZO1-Curved or S2472E-Curved to the S2472E-Intermediate structure (Fig. 2b-e). The spring-gate is switched from an extended to a compressed state, resulting in a change of the central axis-facing F2460 (side chain distance of 9 Å) to Y2464 (side chain distance of 17 Å) (Fig. 2e, f). Compared to the PIEZO1-Curved structure, the volume of the CV, EV and membrane vestibule (MV, between I2466 and the hydrophobic TM-gate residue V2476) of S2472E-Intermediate are increased from ∼1.6 nm^3^ to 3.4 nm^3^, 1.3 nm^3^ to 2.1 nm^3^ and 0.5 nm^3^ to 1.2 nm^3^ respectively, while the inner vestibule (IV, between K2479 and the intracellular constriction neck) remains comparable (Fig. 2g). For the PIEZO1-Flattened structure, while the MV is enlarged, its CV and EV are similar to those of the PIEZO1-Curved (Fig. 2g).

The hydrophilic IV is connected laterally to the intracellular lateral portals and vertically to the constriction neck (Fig. 2c, d). Previous structure-function characterizations^25^ and molecular dynamic simulations^26^ suggest that the lateral portals rather than the constriction neck represent the major ion-permeation routes. Consistent with this, the vertical constriction neck remains closed in the S2472E-Intermediate structure (Fig. 2c, d). The ion-conducting lateral portals are equipped with lateral-plug-gates, which are predicted to be partially unplugged upon channel opening^25^. However, in the S2472E-Intermediate structure, the lateral plug appears not to show apparent structural changes (Extended Data Fig. 6d). The partial flattening of the blade in the S2472E-Intermediate structure might not be sufficient to cause a structural displacement of the lateral-plug-gate. The relatively low resolution of the fully flattened S2472E-Flattened structure has prevented us from analyzing the potential structural change of the lateral-plug-gate. Nevertheless, even in the PIEZO1-Flattened structure with completely flattened blades, the lateral plug remains largely unaltered^6^. Thus, we reason that the lateral-plug-gate could be too dynamic to be captured in the open state. The top portion of the cap remains closed above the residue R2295 position among all the structures (Fig. 2b, c), consistent with mutagenesis studies suggesting that ions might not permeate through the top of the cap^27^. Regardless of the intracellular lateral-plug-gate, the full dilation of the central pore axis comprising the CV, EV and MV suggests that the S2472E-Intermediate structure might represent an open state conformation of the central pore. Furthermore, since the domain containing the lateral-plug-gate of PIEZO1 can be alternatively spliced to produce the widely expressed splicing variant PIEZO1.1^25^, the S2472E-Intermediate structure could represent a fully open state of PIEZO1.1.

### Up to down state of the cap

When positioned in the up-state in the PIEZO1-Curved structure, the Cap^α1^ helix of the cap directly interacts with the extracellular loop linking TM31 to TM32 (EL31-32) in the transmembrane helical unit 8 (THU8) through electrostatic interactions between R1761/R1762 and E2257 (Fig. 3a, b). In the S2472E-Intermediate structure, the down-state position of the cap and partially flattening of the blade leads to disruption of the interaction between Cap^α1^ and EL31-32, and formation of an interaction between Cap^α2^ and the resolved extracellular loop linking TM33 to 34 (EL33-34) in THU9 (Fig. 3c, d). Such a conformational switch might be critical for mechanical activation of PIEZO1. In line with this, deleting the Cap^α1-α2^ or crosslinking between R1761/R1762 and E2257 impaired mechanical activation of PIEZO1^16,28^. Furthermore, we found that deleting a portion of the EL33-34 (K2000-P2017) containing many polar residues (Fig. 3d) abolished mechanical activation of the mutant channel despite its normal expression in the plasma membrane (Fig. 3o and Extended Data Fig. 8a, b). These data support the structural role of EL33-34 in stabilizing the cap in the down state.

### Rotation to open the lateral cap-gate

In the trimeric cap of the PIEZO1-Curved structure, the Cap^β5-β6^-loop (residues R2325-Y2335) and the Cap^β8-β9^ loop (residues R2377-E2383) between neighboring subunits are in close proximity so that the lateral access into the inner cavity of the cap is blocked (Fig. 3e, f). For instance, the side-chain distance between A2328 and P2382 or D2326 and E2383 is ∼4.3 Å and 4.8 Å (Fig. 3e, f), respectively. In the cap of the S2472E-Intermediate structure, the two loops are separated apart so that the side-chain distance between A2328 and P2382 or D2326 and E2383 is increased to ∼16.2 Å and ∼12.8 Å, respectively (Fig. 3g, h). To better appreciate the lateral motion, the cap of the PIEZO1-Curved and S2472E-Intermediate structures is overlayed by aligning at the Cap^α1-α2^ of one subunit, and a top-down view shows that the other two subunits clearly display an anti-clock-wise rotation, as evidenced by the displacement of the central cavity-facing Y2335 for ∼10 Å, widening the lateral access and the central cavity (Fig. 3i). Electrophysiological characterizations have shown that crosslinking A2328 and P2382 prevented mechanical activation of PIEZO1, indicating that conformational flexibility of the lateral access is required for ion conduction^28^. Electrostatics analysis reveals that this access is clustered with negatively charged residues including D2326, E2334, E2338, E2383 and DEEED (2393-2397) (Fig. 3g, h), which might collectively help to enrich cations over anions for permeation. In line with this, mutating DEEED (2393-2397) to AAAAA resulted in the mutant channel with increased permeability of chloride^29^. These structural-functional analyses led us to propose that the Cap^β5-β6^-loop and the Cap^β8-β9^-loop between neighboring subunits might form lateral cap-gates, and accordingly we term the two loops as cap-gate-loop 1 and 2 (Fig. 3f, k).

### Extended to compressed state of the spring-linker

The cap is linked to the pore-lining IH through the linker region of S2450-G2465 following the Cap^β11^ (Fig. 3j, k and Extended Data Fig. 6e). Given that the conformational change of this linker resembles the change of a spring from an extended to a compressed state and its critical role in mechanically gating PIEZO1, we name this linker the spring-linker (Fig. 3j, k). On the basis of the PIEZO1-Curved, PIEZO1-Flattened and S2472E-Curved structures, the spring-linker adopts an extended state to enclose the hydrophobic EV (Fig. 2c-f and Fig. 3e). In the S2472E-Intermediate structure, the whole spring-linker is fully resolved and the lower portion is folded into a short helix in parallel to the membrane plane (Fig. 3j, k and Extended Data Fig. 6e). D2451 in the upper portion of the loop (S2450-P2456) is close to R2377 in the cap-gate loop 2 with a side chain distance of ∼2.7 Å, while K2452 is ∼3-4 Å proximity to E2279 and D2280 located in the turning loop connecting Cap^α2^ to Cap^β4^ (Fig. 3k). Mutating both D2451 and K2452 or E2279 and D2280 to alanine (D2451A/K2452A or E2279A/D2280A) totally abolished mechanical activation without affecting the expression on plasma membrane (Fig. 3o and Extended Data Fig. 8). Single residue mutations of D2451 (D2451A, D2451K and D2451R) and R2377 (R2377D), but not K2452 (K2452A) abolished mechanical activation (Fig. 3o). Interestingly, the loss of mechanically activated current in the D2451R and R2377D mutants was partially rescued in the D2451R/R2377D double mutant (Fig. 3o). These data suggest that the electrostatic interaction between D2451 and R2377 might play a critical role in controlling the cap-gate.

The lower helical portion of the spring-linker is laterally pulled away from the central axis, resulting in the hydrophilic Y2464 (∼13 Å displacement) rather than the hydrophobic F2460 (∼16 Å displacement) becoming the EV-facing residue and drastic expansion of the EV (Fig. 2f and Fig. 3m). In line with the drastic conformational change, when the helical portion (P2455-Y2464) is deleted, the ΔSpring-linker mutant channel completely loses mechanical activation (Fig. 3o), which might be in part due to its reduction in the expression (Extended Data Fig. 8). Our previous studies have found that mutants including L2461C, A2462C, G2465C have drastically reduced mechanically evoked currents, and Y2464C is able to be covalently labeled by MTSES to block the current^29^. Intriguingly, azobenzene-based photoswitch covalently tethered to Y2464C was shown to confer light-induced gating of the mutant PIEZO1 channel^30^. Taken together, these structural and functional characterizations demonstrate that the spring-linker serves as a key structural element in the mechanical gating of PIEZO1.

### Lateral pulling of the IH through hydrophobic interactions

The OH is linked to the Cap^β1^ through the OH-Cap-loop (S2215-P2223) (Extended Data Fig. 6e). In the S2472E-Intermediate structure, the displacement of the cap leads to a concurrent downward and outward movement of the OH-Cap-loop and a lateral bending of the outer portion of OH for ∼13 Å, resulting in an apparent kink at the residue P2206 (Fig. 3k, l). Notably, the outer portion of OH consists of the patch of hydrophobic residues from I2202 to L2212 (IIWFPLLFMSL), which forms extensive hydrophobic interactions with the outer portion of IH (residues I2466, L2469, V2474, V2477) (Fig. 3d). The lateral motion of the spring-linker and the OH-Cap-loop and bending of the outer portion of OH might allow the hydrophobic interaction to laterally pull the IH away from the central axis (Fig. 3n and Supplementary Video 3, 4), and consequently dilate the hydrophobic pore. Compared to the curved structure, the IH of the S2472E-intermediate structure has a lateral movement at the upper residue I2466 for ∼10 Å and the lower residue K2479 for ∼4 Å (Fig. 3m).

### Returning the cap to the up state and closing the cap-gate

Compared to the S2472E-Intermediate structure, the cap and spring-linker of the PIEZO1-Flattened structure is released back to the up- and extended-state (Fig. 2a-c and Fig. 4a), which is similar to that of the PIEZO1-Curved and S2472E-Curved structures, resulting in closure of the extracellular cap-gate and spring-gate (Fig. 4b-d). Furthermore, the cap, TM blade and IH of the PIEZO1-Flattened show a clock-wise rotation in relative to that of the S2472E-intermediate (Fig. 4a), a reversed mode of action observed from the curved state to the S2472E-Intermediate state (Extended Data Fig. 6c). Compared to the S2472E-Intermediate structure, the three blades of the PIEZO1-Flattened were contracted but extended peripherally, resulting in the V650 has a shift of ∼44 Å (Fig. 4a). The maintenance of the blade in the flattened configuration might hold the TM-gate still in a relatively dilated state with enlarged MV (Fig. 2b-d, g).

**Figure 4.**
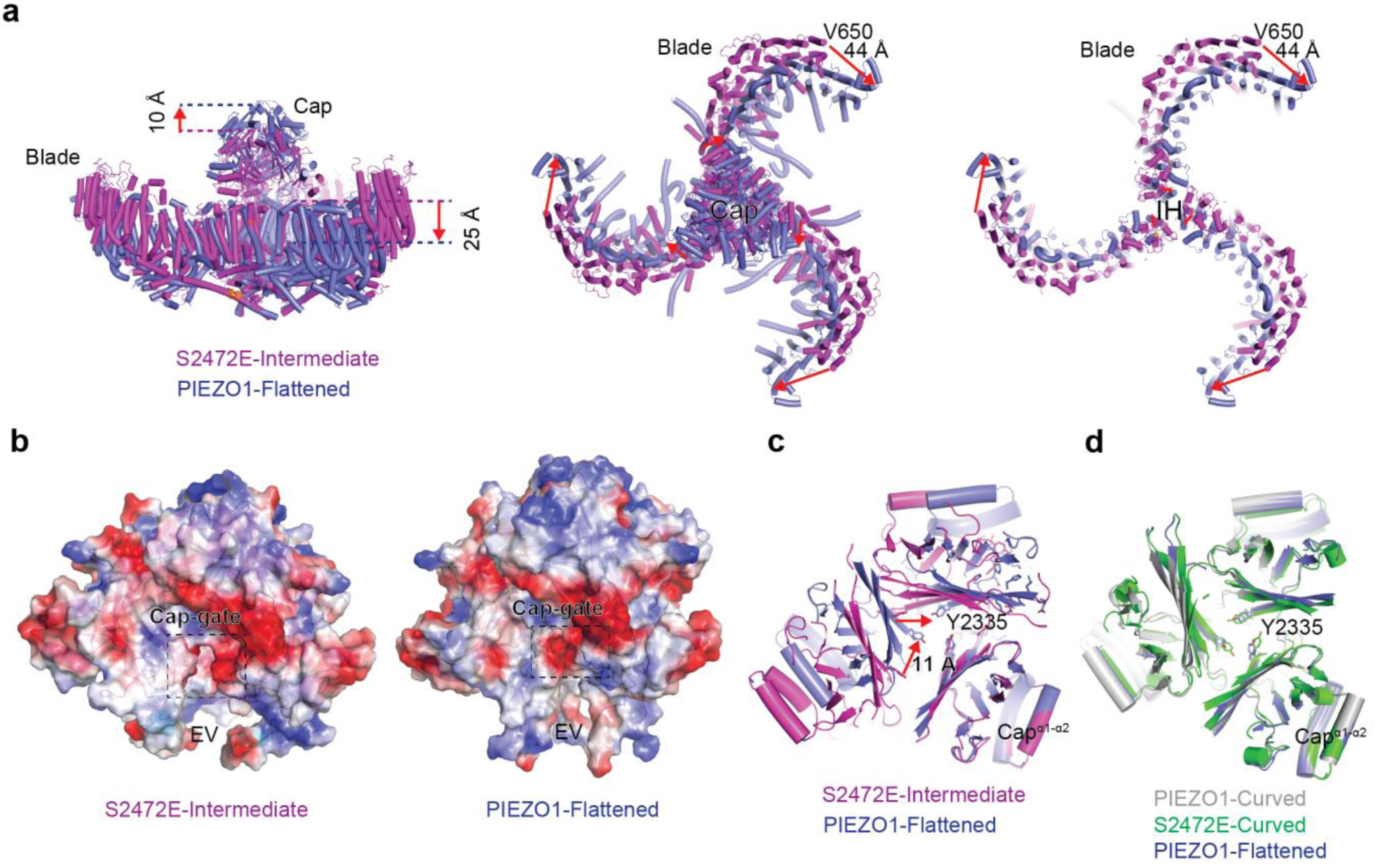
| Conformational changes from S2472E-Intermediate to PIEZO1-Flattened. **a**, The overlaid cartoon models of S2472E-Intermediate and PIEZO1-Flattened structures viewed in side (left), top (middle), or top with the extracellular cap being omitted (right) to clearly display the pore-lining IH and blade TMs. Red arrows indicate the conformational motion from the S2472E-Intermediate to the PIEZO1-Flattened structure. **b**, Electrostatic surface potential and model of the cap of the S2472E-Intermediate and PIEZO1-Flattened. The dash box outlines the cap-gate. **c** and **d**, Superimposed top-down view zoomed at the cap-gate residue Y2335 of the indicated structures aligned at the Cap^α1-α2^ of one subunit. Red arrows indicate the displacement of the Y2335 residues in the other two subunits.

### MD simulations of ion permeation

The collective motion of up-to-down and opening of the cap-gate, the extension-to-compression of the spring-gate and the lateral dilation of the TM-gate lead to a continuation and expansion of the CV, EV, and MV in the S2472E-Intermediate structure (Fig. 2). Thus, we hypothesize that the structure might represent an open state of the central pore to allow ion permeation sequentially through the lateral cap-gate, the spring-gate, and the TM-gate. To validate this, we have conducted coarse-grained MD simulations of the truncated structure containing the central pore module (residues 1956-2547), the beam (residues 1315-1364/1365) and the intracellular lateral-plug-gate (residues 1401-1421) at four different transmembrane potentials of 0 V, -0.1 V, -0.25 V, and -0.5 V, respectively. In the PIEZO1-Curved structure, while Na^+^ and water molecules were able to access the EV (Fig. 5a and Extended Data Fig. 9a), only a few Na^+^ ions were found to be occasionally trapped in the CV, and a portion could reach the MV but none permeated through the hydrophobic TM-gate residue V2476 to arrive at the IV even at -0.5 V (Fig. 5a, c, Extended Data Fig. 9b and Supplementary Video 5), suggesting a non-conducting state. The near-zero water density around the hydrophobic constriction site in the PIEZO1-Curved structure is consistent with previous MD simulations studies^31^. In contrast, in the S2472E-Intermediate structure, we observed Na^+^ permeation in a transmembrane potential-dependent manner (Fig. 5a, c). The larger the voltage, the more Na^+^ permeated to the IV (Fig. 5a, c and Extended Data Fig. 9b). As shown in the Supplementary Video 6 and the series of snapshots at -0.5 V (Fig. 5b), Na^+^ readily accessed the CV through the three-lateral cap-gates, then flowed to the EV and MV (Fig. 5a-c). Quantitative analyses reveal that a comparable number of Na^+^ accumulatively accessed the CV, EV and MV during the simulation period (for instance, 62, 81, 85 at -0.5 V or 27, 23, 24 at -0.1 V, respectively), but transited through the hydrophobic V2476 position to the IV in a voltage- and time-dependent manner (Fig. 5a-d and Extended Data Fig. 9b, c). The ion permeation rate from MV to IV is ∼39%, 42%, 53% and 78% at 0 V, 0.1V, 0.25 V and 0.5 V, respectively (Fig. 5c and Extended Data Fig. 9b). On the basis of the number of Na^+^ ions permeated through the TM pore at different voltages, we calculated the single-channel current and slope-conductance of ∼20 pS (Fig. 5e), which was close to the electrophysiologically measured conductance of ∼30 pS^1^. Taken together, these data demonstrate that the S2472E-Intermediate structure might represent a central pore-opening state of PIEZO1, in which V2476 serves as a key hydrophobic gate to control ion permeation through the TM pore.

**Figure 5.**
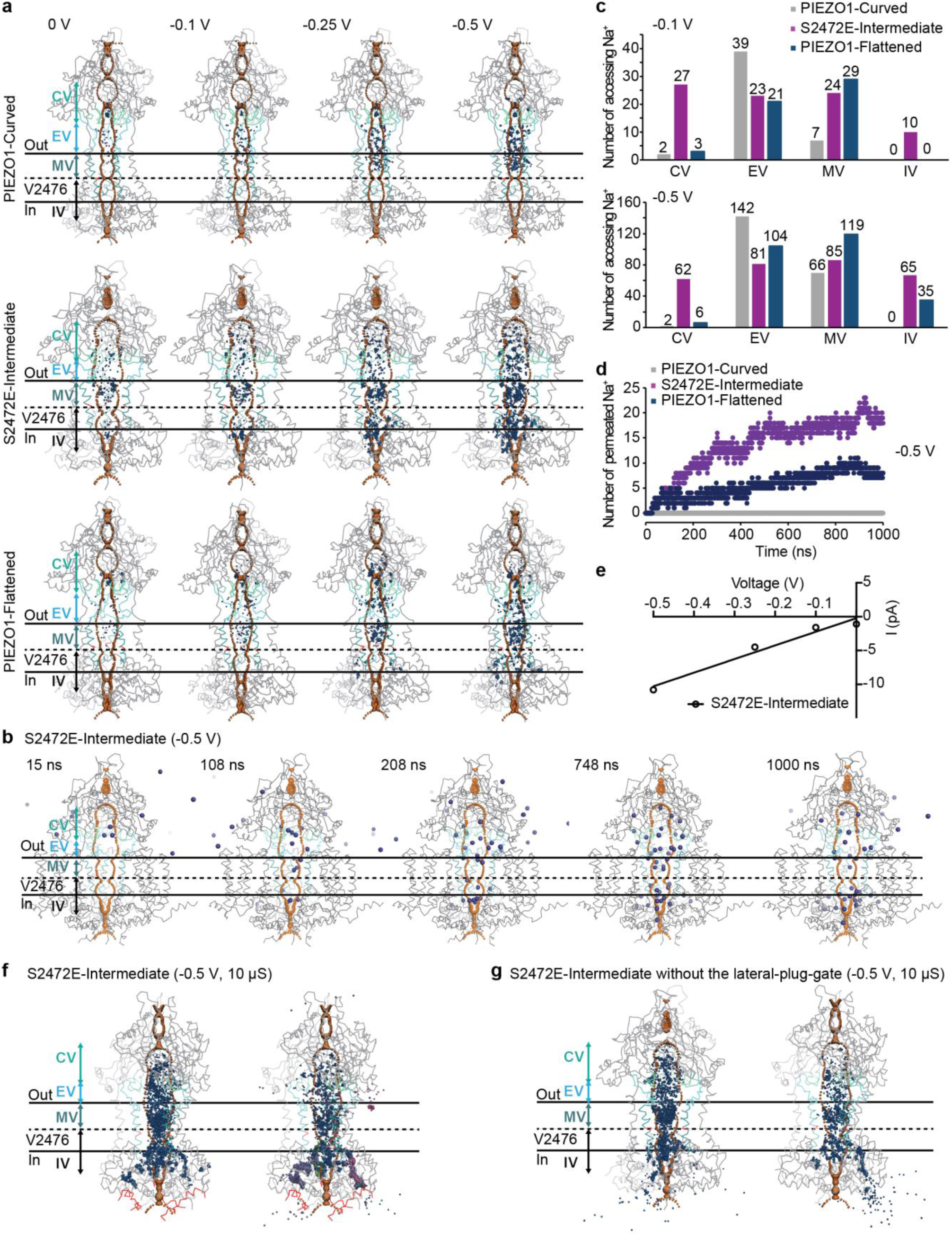
| Molecular dynamic simulations of ion permeation. **a**, Molecular dynamic simulations of the Na^+^ assessing to the indicated CV, EV, MV and IV of the indicated structures. The central pore boundary is illustrated with orange dots generated from HOLE, and the accessing Na^+^ ions are represented by the blue dots. The location of the hydrophobic gate residue V2476 is shown in red and indicated by the black dash line. **b**, A series of snapshots of Na^+^ permeation through the CV, EV, MV and IV of S2472-Intermediate structure simulated at -0.5 V. **c**, The accumulated number of Na^+^ ions accessing the indicated cavities of the structures at -0.1 V or -0.5 V. **d**, Time-dependent permeation of Na^+^ to the IV of the PIEZO1-Curved (grey), S2472E-Intermediate (purple) and PIEZO1-Flattened (dark blue) at -0.5 V. **e**, Current-voltage curve of Na^+^ permeated through the TM pore of the S2472E-Intermediate structure. **f**, MD simulations of the S2472E-Intermediate structure at -0.5 V for 10 µs show the tracked trajectories of all Na^+^ ions during the last 1 µs of the simulation that permeated through the TM pore into the IV and flowed along the intracellular lateral portal (left panel), or the 5 Na^+^ ions (represented in different colors) in 10 µs that permeated through the lateral-plug-gate (shown in red) (right panel). **g**, MD simulations of the S2472E-Intermediate structure without the lateral-plug-gate at -0.5 V for 10 µs show the tracked trajectories of all Na^+^ ions during the last µs of the simulation that permeated through the TM pore into the IV and flowed along the intracellular lateral portal (left panel), or the 37 Na^+^ ions in the last 1 µs permeated through a single lateral portal and enter into the intracellular space at the deleted lateral-plug-gate location (right panel).

To examine whether the IV-residing Na^+^ might flow along the lateral portal and enter into the intracellular side through the lateral-plug-gate, we prolonged the simulations on the S2472E-Intermediate structure to 10 µs at 0.5 V. The tracked trajectories of all IV-reaching Na^+^ ions show that a considerable number of Na^+^ were observed in the lateral portal, but only 5 ions managed to traverse the lateral-plug-gate (Fig. 5f), consistent with the hypothesis that the lateral-plug-gate needs to be unplugged to allow ion permeation. To further test this, we conducted similar simulations on the S2472E-Intermediate structure with the lateral plug domain removed, which might resemble a fully open state of the ion conduction pathway. In the last microsecond of the 10 µs simulations, a total of 72 Na^+^ ions permeated through the TM pore to reach the IV. Remarkably, 37 Na^+^ ions flowed along a single lateral portal and entered into the intracellular space via the removed lateral-plug position (Fig. 5g). The estimated conductance of the IV region is 23 pS, similar to the simulated slope conductance (Fig. 5e). Interestingly, the estimated conductance through a single lateral portal without the lateral-plug-gate is 12 pS, which is close to one third of the single-channel conductance of 44 pS of the PIEZO1.1 isoform that lacks the lateral-plug-gate^25^. Notably, no Na^+^ ions were observed to permeate through the central constriction neck under all the simulated conditions. Thus, consistent with previous MD simulations^26^ and mutagenesis and electrophysiological characterizations^25^, our simulation results support that the lateral portal and lateral-plug-gate serve as the intracellular ion-conducting pathway of PIEZO1.

To test the ion conductivity of the PIEZO1-Flattened structure, we conducted MD simulations and found that the access of Na^+^ to the EV and MV of PIEZO1-Flattened was comparable to that of S2472E, but the permeation rate through the hydrophobic gate of V2476 is much reduced (∼10%, 0%, 16% and 29% at 0 V, -0.1 V, -0.25 V and -0.5 V) (Fig. 5a, c, Extended Data Fig. 9b, c). Consistent with closed cap-gate, only few Na^+^ were trapped in the CV even at -0.5 V (Fig. 5a, c, Extended Data Fig. 9b, c, and Supplementary Video 7). These analyses suggest that the PIEZO1-Flattened structure might represent an inactivated state with closed extracellular cap-gates and extended spring-linkers. In line with this, both the cap-gate-loop 1 and 2 are functionally involved in determining the distinct inactivation kinetics of PIEZO1 and PIEZO2^28^.

## Discussion

Combining analysis of the series of PIEZO1 and S2472E structures, mutagenesis and electrophysiological characterizations and MD simulations, here we assign the close, intermediate, open and inactivated states of PIEZO1 with characteristic structural features (Fig. 6). In the closed state represented by the PIEZO1-Curved and S2472E-Curved structure, PIEZO1 adopts an intrinsically curved trimeric structure with a measured middle-plane radius of ∼10-13 nm, in which the blades are curved, the cap is in the up-state and all the gates are closed (Fig. 6a). On the basis of the S2472E-Intermediate structure with the open central pore and the existence of the S2472E-Flattened structure whose blades are fully flattened and the cap is in the down-state, we propose that the S2472E-Intermediate structure represents an intermediate open state in which lowering and rotating the cap, partially flattening the blade and levering the beam lead to opening of the cap-, spring- and TM-gates to allow ion permeation through the central pore, while the lateral-plug-gate remains closed (Fig. 6b, Supplementary Movie 4-6). MD simulations of the S2472E-Intermediate structure without the lateral-plug domain revealed ion permeation into the cytosolic side, supporting unplug of the lateral-plug-gate in the full open state of PIEZO1 (Fig. 6c). While it has been proposed that the peripheral blades serve as mechanosensing domains to open the central pore via a periphery-to-center transduction mode^32^, the S2472E-Intermediate structure suggests a reversed center-to-periphery mode, demonstrating the reciprocal force transduction between the peripheral blade-beam and the central cap and pore module. The open state is transited to the inactivated state represented by the PIEZO1-Flattened structure through simply releasing the cap and spring-linker back to the up- and extended state for closure (Fig. 6d). Returning the flattened blades to the curved state is expected to further close the TM-gate and lateral-plug-gate, setting the channel back to the closed state.

**Figure 6.**
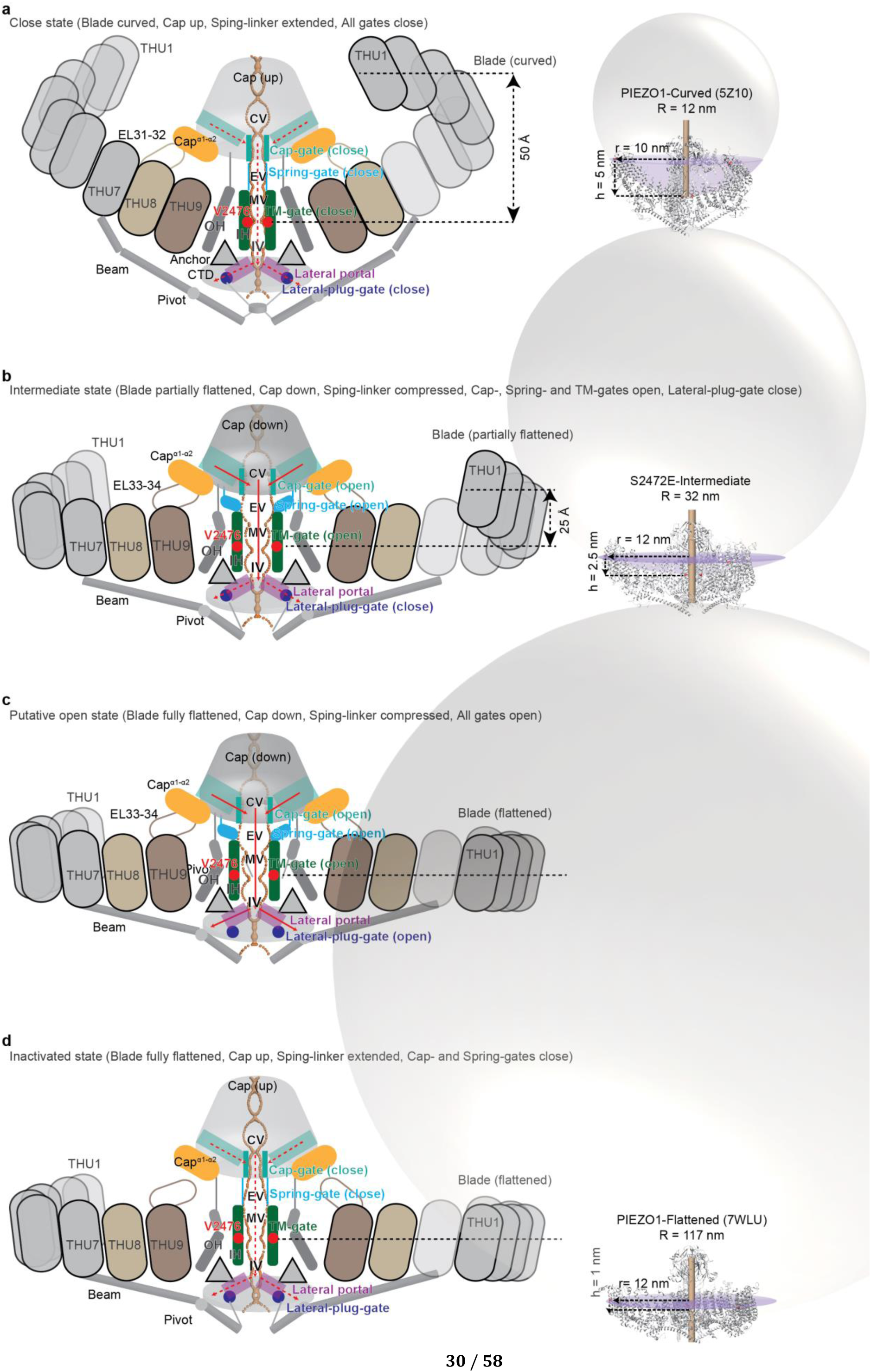
Schematic of the structural choreography and curvature-based gating of PIEZO1. Cartoon model representation of the close (**a**), intermediate (**b**), open (**c**), and inactivated-state (**d**) of PIEZO1. Dashed and solid red arrows indicate the blocked and opened ion permeation pathway, respectively. THU represents transmembrane helical unit containing 4 TMs. The red dots indicate the hydrophobic TM-gate residue V2476. Key components are labeled. On the basis of the S2472E-Intermediate and S2472E-Flattened structures and MD simulations, the putative open state model (**c**) is proposed, in which the partial unplug of the lateral-plug-gates remains to be structurally determined. The curvature radius (R) of PIEZO1-Curved (PDB: 5Z10), S2472E-Intermediate and PIEZO1-Flattened (PDB: 7WLU) are calculated and illustrated with the spherical ball, the purple area represents the bowl area.

PIEZO channels contain multiple gates including the extracellular cap-gate and spring-gate, the hydrophobic TM-gate, and the intracellular lateral-plug-gate that can be alternatively spliced. The multiple gate system might safeguard PIEZO channels to serve as specialized sensors to mechanical force, but not to other chemical or physical stimuli such as temperature. The unique structural design might allow the coordinated conformational changes of the key mechanogating components (blade, beam, cap, and spring-linker) to simultaneously open all the gates specifically in response to changes in mechanical force with an estimated half-maximal activation tension of ∼1.4-1.9 mN/m^6,33^, but not to other stimuli. At the meantime, the multiple gate system might provide a great flexibility for regulating the functional state of PIEZO channels to meet their wide variety of cellular functions. On the basis of the proposed transitional states from close to intermediate, open and inactivation, it is intriguing that the S2472E-Intermediate structure has ∼25 Å flattening of its blades and associated opening of the cap-, spring- and TM-gates, while the intracellular lateral-plug-gates remain closed (Fig. 6b). A further flattening of the blade for another 25 Å might be needed to partially unplug the lateral-plug-gates so that the whole ion permeating pathway is fully open (Fig. 6c). Such a correlation appears to be remarkable as it might suggest that the flattening of the blade serves as a scalable control of the cap-gate/spring-gate/TM-gate and the lateral-plug-gate. In line with this hypothesis, the widely expressed PIEZO1.1 splicing variant lacks the intracellular lateral-plug-gate, but remains to be responsive to mechanical forces with about two-fold of enhanced mechanosensitivity^25^. The S2472E-Intermediate structure might represent a full open-state of the PIEZO1.1 isoform, which only requires half of the flattening of the blade for opening. Furthermore, since closing any of the gates might prevent ion permeation, inactivation might occur not only at the extracellular cap- and spring-gates, but also at the TM-gate. In line with this, mutating L2475 and V2476 to hydrophilic serine drastically slowed inactivation, leading to proposal of the hydrophobic inactivation gate^22^. Thus, we propose that PIEZO channels might adopt varied inactivation states to account for their distinct inactivation kinetics under physiological and pathophysiological conditions, which might require additional structural validation.

The measured middle-plane curvature radii (R) of the PIEZO1-Curved (PDB: 7WLT, 5Z10 and 6B3R), S2472E-Curved, S2472E-Intermediate and PIEZO1-Flattened (PDB: 7WLU) structures are ∼10-12 nm, 14 nm, 32 nm, 117 nm, respectively. The higher R value of S2472E-Curved than that of PIEZO1-Curved could be a corelative indication of the higher basal activities observed in S2472E-expressing cells (Fig. 1 and Extended Data Fig. 1d, e). Mathematical models have predicted that the PIEZO dome (PIEZO channel together with the residing lipid membranes) adopts a resting state with an R value of ∼42 nm, on the basis of considering lipids devoid of any tendency towards spontaneous curvature such as phosphatidylcholine (PC), which form planar but not curved bilayers^34,35^. Nanoscopic fluorescence imaging of PIEZO1 expressed on the plasma membrane has measured the inter-blade distance at the distal end (d) of PIEZO1 at resting state to be ∼25 nm^36^, which corresponds to the in-plane radius (r = d/√3) of ∼14 nm according to the calculation method shown in Extended Data Fig. 7b. If assuming the height of the blade (h) varied at 5 nm, 4 nm, 3 nm and 2.5 nm (similar to the S2472E-intermediate state), the according R value is 23 nm, 28 nm, 36 nm and 43 nm, respectively. If PIEZO1 at resting state in cell membrane adopts the predicted curvature radius of ∼42 nm, its central pore is expected to be at least as open as the S2472E-Intermediate state with an R value of ∼32 nm. However, this prediction contradicts the functional observation that S2472E had higher basal activities than PIEZO1 (Fig. 1 and Extended Data Fig. 1d, e). Therefore, we reason it is more likely that PIEZO1 in cell membrane at resting state might adopt a curvature radius between 23 to 32 nm with only slight flattening. These analyses suggest that PIEZO1 can adopt varied degree of curvatures, leading to a transition from the intrinsically curved state of ∼10-12 nm radius, to the prone-open S2472E-Curved of ∼14 nm radius, to less curved state of ∼23-32 nm radius under native membrane composition, to a partially flattened and partial open state of ∼32 nm radius, until reaching the fully flattened and open state (Fig. 6). Thus, the S2472E-Intermediate structure with verified open central pore is highly valuable not only for revealing the intrinsic structural choreography of PIEZO1, but also for understanding the variable conformational states of PIEZO1 in native cell membrane. In summary, our studies have provided direct structural evidence in supporting the curvature-based mechanogating mechanisms of PIEZO1 and raised the intriguing hypothesis that PIEZO1 might utilize scalable deformation for precise mechanosensing.

## Methods

### Molecular Cloning

All PIEZO1 mutant constructs including S2472E, R2482H, ΔEL33-34 (K2000-P2017), ΔSpring-linker (P2455-Y2464), E2279A/D2280A, D2451A/K2452A, K2452A, D2451A, D2451K, D2451R, R2377D, D2451R/R2377D were sub-cloned using the One Step Cloning Kit (Vazyme Biotech) according to the instruction manual and as previously described^29^. The WT and mutated PIEZO1 cDNAs were cloned into the pcDNA3.1 expression vector. For protein purification, the PIEZO1-S2472E cDNA was linked to a C-terminal glutathione S-transferase (GST) tag with a precision protease cleavage site in between, followed by the GFP-coding sequence driven by an internal ribosome entry site (IRES) (for monitoring the efficiency of transient transfection and protein expression). For electrophysiology recordings, the PIEZO1-WT and mutated cDNAs were fused directly with the mRuby2 fluorescent protein-encoding sequence for accurately reflecting the expression and location of the PIEZO1 proteins.

### Whole-cell electrophysiology

All the electrophysiology tests were performed on PIEZO1-KO-HEK293T cells in which the endogenous PIEZO1 gene is deleted. Cells were transfected with PIEZO1 or mutant fused with GFP or mRuby using Lipofectamine 3000, and GFP or mRuby fluorescent signals were checked 24 h later for monitoring transfection and expression efficiency. Cells were briefly digested with diluted 0.05% trypsin (1:20) and sparsely re-plated in poly-D-lysine-coated coverslips and incubated around 2 h for recording prior to patch clamp. Patch-clamp experiments with HEKA EPC10 were based on those previously described^16,17^. For whole-cell patch-clamp recordings, the recording electrodes had a resistance of 2–5 MΩ when filled with an internal solution composed of 133 mM CsCl, 1 mM CaCl_2_, 1 mM MgCl_2_, 5 mM EGTA, 10 mM HEPES (pH 7.3 with CsOH), 4 mM MgATP and 0.4 mM Na2GTP. The extracellular solution was composed of 133 mM NaCl, 3 mM KCl, 2.5 mM CaCl_2_, 1 mM MgCl_2_, 10 mM HEPES (pH 7.3 with NaOH) and 10 mM glucose. All experiments were carried out at room temperature. The currents were sampled at 20 kHz and filtered at 2 kHz using Patchmaster software. Leak currents before mechanical stimulation were subtracted off-line from the current traces. Mechanical stimulation was delivered to the cell during the patch-clamp recording at an angle of 80° using a fire-polished glass pipette (tip diameter 3–4 μm) as previously described^1^. The downward movement of the probe towards the cell was driven by a Clampex-controlled piezoelectric crystal micro-stage (E625 LVPZT Controller/Amplifier; Physik Instrument). The probe had a velocity of 1 μm ms^−1^ during the downward and upward motion, and the stimulus was maintained for 150 ms. A series of mechanical steps in 1 μm increments was applied every 8 s and the currents were recorded at a holding potential of −80 mV.

### Single-cell Fura-2 Ca^2+^ imaging

HEK293T cells grown on 8-mm round glass coverslips coated with poly-D-lysine and placed in 48-well plates were transfected with either PIEZO1 or S2472E, and then were subjected to Fura-2 Ca^2+^ imaging 36 h post transfection according to the previously described protocol^37,38^. The confluent cells were washed with buffer containing 1 × HBSS (with1.3 mM Ca^2+^ and 10 mM HEPES (pH 7.2 with NaOH), then incubated with 2.5 μM Fura-2-AM (Molecular Probes) and 0.05% Pluronic F-127 (Life technologies) for 30 min at room temperature. After washing with the buffer, the coverslips were mounted into an inverted Nikon-Tie microscopy equipped with a CoolSNAP charge-coupled devise (CCD) camera and Lambda XL light box (Sutter Instrument). GFP positive and -negative cells were selected for measurement of the 340/380 ratio with a 20× objective (numerical aperture N.A. = 0.75) using the MetaFluor Fluorescence Ratio Imaging software (Molecular Device). A stock solution of Yoda1 at 30 mM was solubilized in DMSO and diluted to a final concentration of 5 μM. The compound solutions were perfused into the chamber and cells via a multichannel perfusion system (MPS-2, World Precision Instruments).

### Cell viability assay

HEK293T cells per well (100 μ media) were seeded in 96-well plates. Typically, cells were transfected with the mixture of 500 ng PEI and 200 ng plasmids as indicated, and the cell viability was determined using CellTiter-Glo viability assay kit (Promega) at different time points according to the manufacturer’s protocols. Briefly, 100 μL CellTiter-Glo Reagent which is equal to the volume of cell culture medium was added into each well. After 10 mins incubation on a room temperature shaker, the luminescent signals were recorded by a microplate reader (EnVision, Perkin). The luminescence from reagent-lysed cells is proportional to the ATP content and, thus, is proportional to the number of live cells that are present. Each independent experiment was performed in quadruplicate and repeated twice.

### Expression and purification of the PIEZO1-S2472E mutant protein

The expression of the PIEZO1-S2472E-mRuby2 proteins was monitored by taking fluorescent and bright field images every 6 hours after transfection. The purification procedure was adapted from previously described methods^6,17^. To overcome the obstacle that overexpression of S2472E in HEK293T cells led to cell death and consequently insufficient S2472E-expressing cells for protein purification, we have optimized the transfection conditions after cells reaching 95-100% confluency to obtain acceptable cell viability with reasonable expression level of the S2472E mutant. In this manner, we collected and processed transfected cells from 140 150 mm-petri dishes for each prep of protein purification. Briefly, the PIEZO1-S2472E-GST (Glutathione S-transferase) fusion protein, with a precision protease cleavage site in between, was expressed 24 hours post-transfection with the expression vector. Green fluorescent protein (GFP)-coding sequence, driven by an internal ribosome entry site (IRES), was incorporated to monitor the efficiency of transient transfection and protein expression. Cells were collected and homogenized by lysis buffer, containing 25 mM Na-PIPES, pH 7.3, 150 mM NaCl, 0.5% (wt/vol) L-a-phosphatidylcholine (Avanti), 3 mM DTT, a cocktail of protease inhibitors (Roche), and a mixture of detergents (1% (wt/vol) CHAPS, 0.1% C12E10, 0.05% GDN 0.05%) for 2 hours. After centrifugation at 140,000g for 30 min, the supernatant was collected and incubated with glutathione– sepharose beads at 4 °C for 6 hours. The PIEZO1-S2472E mutant protein was cleaved from the resin by 16 hour-incubation with 0.1 mg/ml prescission protease, and further purified using size-exclusion chromatography (Superpose-6 10/300 GL, GE Healthcare) in the buffer containing 25 mM Na-PIPES, pH 7.3, 150 mM NaCl, 3 mM DTT, and 0.02% GDN. Peak fractions between 9.0 ml and 10.0 ml containing well-dispersed trimeric PIEZO1-S2472E proteins were collected for further assays.

### Cell surface biotinylation and western blotting

Cell surface biotinylation and western blotting were carried out as previously described. In brief, cultured HEK293T were transfected with the following plasmid: ΔEL33-34, ΔSpring-linker, D2451A/K2452A, E2279A/D2280A. After washing by DPBS with Ca^2+^ and Mg^2+^, cells were incubated with 0.5 mg/mL EZ-Link NHS-LC-Biotin (Thermo Fisher Scientific) for 45min, then quenched by 100 mM glycine solution. The cells were then lysed and incubated with streptavidin magnetic beads overnight. The input sample and washed beads sample were subjected to 8% SDS-PAGE gels and transferred to 0.45 μm PVDF membranes (Millipore), the membrane was blocked by 5% Nonfat dry milk (BIO-RAD) in TBST buffer (TBS buffer with 0.1% Tween-20) and incubated with the primary antibodies overnight. After washing with TBST buffer, the membrane was incubated with the peroxidase-conjugated secondary antibody at room temperature for 1 h, followed by washing and detection using an enhanced chemiluminescence detection kit (Thermo Fisher Scientific). The antibodies used for western blotting included rabbit anti-PIEZO1 (custom generated, 1:2,000) and rabbit anti-β-actin (Cell Signaling Technology, 1:3,000).

### Negative-staining electron microscopy

4 μl freshly prepared sample (about 0.04 mg ml^-1^) was loaded to a glow-discharged (PELCO) carbon-coated copper grid (300 mesh, Zhongjingkeyi Technology) and incubated for 1 minute. The excess sample liquid was removed by filter paper^39^. After 15 s, the excess buffer was removed with filter paper, and then the grid was stained by a droplet of 2% uranyl acetate for 20 s. The staining process was repeated twice and the third staining process was carried out additionally 30 s. Finally, the excess staining buffer was removed, and the grid was dried at room temperature. The prepared girds were observed on a T12 microscope (FEI) operating at 120 kV, using a 4k x 4k CCD camera (UltraScan, Gatan) at a nominal magnification of 49,000 with a calibrated pixel size of 2.3 Å.

### Sample preparation and cryo-electron microscopy data acquisition

4 μl of freshly purified mPIEZO1-S2472E sample (around 3 mg ml^-1^) each time was sequentially blotted 3 times to 300-mesh holey carbon Au R1.2/1.3 grids treated with 25s of glow discharge (Quantifoil, Micro Tools GmbH). Fluorinated Fos-choline-8 (0.65 mM; Hampton) was added to the freshly concentrated sample immediately before freezing. The grids were then blotted and plunged into liquid ethane using an FEI Mark IV Vitrobot operated at 8 °C and 100% humidity.

The grids were transferred to a 300 kV Titan Krios (FEI) electron microscope equipped with a Cs corrector and GIF Quantum energy filter (slit width 20 eV). Images were recorded by a post-GIF K3 Summit direct electron detector (Gatan) working at the super-resolution mode. Data acquisition was performed using AutoEMation II with a nominal magnification of 64,000×, which yields a super-resolution pixel size of 0.549 Å on the image plane, and with defocus ranging from -1.5 to -2.0 μm. The total exposure time was set to 2.56s with 0.08s per frame yielding a 32-frame stack. The total dose was approximately 50 *e*-per Å^2^ for each micrograph.

### Image processing

A simplified flowchart of the procedure for image processing is presented in Extended Data Fig. 3 and 5. The image processing of S2472E-Intermediate structure is taken for example, while the image processing of S2472E-Curved structure is similar. Three batches of cryo-EM data containing 7,723, 6,945 and 10,658 micrograph stacks were processed in the first few steps. The stacks were motion-corrected and dose-weighted using MotionCor2^40^ with 2 × 2 binning, resulting in a pixel size of 1.0979 Å. After whole-image CTF estimation using CTFFIND4.1^41^, 7,022, 6,022 and 9,156 good micrographs, respectively, were manually selected. A total of 1,544,600, 1,384,800 and 2,131,100 particles were auto-picked using RELION-3.1^42^. After several rounds of 2D classification, 68,761, 79,192 and 58,761 particles which showed flatter blades were selected. Then 521,038 particles were auto-picked using maps generated by three datasets for templates. After multi-reference 3D classification (K=4), 144,704 particles were selected. The PIEZO1 electron microscopy map (EMD-6865), low-pass-filtered to 60 Å, was used as an initial model. Each particle was re-centered and subjected to Gctf^43^ to refine the local defocus parameters. The well-centered particles with more accurate defocus parameters were re-extracted from the motion-corrected and dose-weighted micrographs and subjected to 3D refinement, which resulted in electron density maps at 4.00 Å resolution. All 3D refinement and 3D classification above were performed with *C*_3_ symmetry.

As the distal blades were absent from the overall 4.00 Å map, focused refinements were performed. The subroutine of Relion_particle_symmetry_expand in RELION-3.1 was used to expand 144,704 particles with *C*_3_ symmetry. Then 434,112 blade particles were obtained using the subtraction of two adjacent blades in raw images. These blade particles were subjected to 3D refinement, resulting in the final focused map of distal blade at 3.74 Å resolution.

In the processing of S2472E-Curved structure, as the cap was absent, a focused refinement of the central region of the 162,903 particles was performed by subtracting the projections of most parts of three surrounding blades. *C*_3_ symmetry was imposed during the refinement with a local mask and resulted in a focused map at 4.54 Å resolution. The two final focused maps were fit together according to their overlapping area using Chimera^44^, then combined using PHENIX^45^ combine_focused_maps. Three copies of the combined map were further combined into the final density map. The reported resolutions are based on the gold-standard Fourier shell correlation 0.143 criterion^46^. All density maps were sharpened by applying a negative *B*-factor that was estimated using automated procedures^47^. Variations in the local resolution were estimated using Resmap^48^.

The S2472E-Flattened structure was discovered through 3D classification of S2472E-Intermediate structure’s image processing as shown in Extended Data Fig. 5. A total of 78,112 particles selected from the initial 3D class were pooled for further processing. Two parallel 3D classifications were performed with two and three classes. Particles from each class were subjected to 3D refinement separately, resulting in two maps at 9.14 Å resolution and 8.66 Å resolution. After a further round of 3D classifications and 3D refinements separately, three density maps at 7.78 Å resolution, 7.95 Å resolution and 8.23 Å resolution were obtained. The duplicated particles derived from three 3D refinements were removed during data merging. After 3D classification(K=4), 3D refinement and post-processing, this dataset yielded a map of S2472E-Flattened at 9.57 Å resolution using cryoSPARC^49^.

### Model building and structure refinement

For the S2472E-Curved structure, the previously determined cryo-EM structures of PIEZO1 (PDB: 5Z10) were aligned to the cryo-EM map by Chimera, and used as the initial models. Manual fitting was performed using COOT^50^. Then the model was refined in real space using PHENIX with secondary structure and non-crystallographic symmetry (NCS) restraints. Lipid-like densities appeared in the density maps were fitted by trial with candidate lipids. For S2472E-Intermediate structure, the PIEZO1 model was used as the starting point, and was split into several parts to be fitted into the map by Chimera. The rest process of model building was similar to the S2472E-Curved structure The final atomic model was validated using MolProbity^51^.

### Measurement of the curvature radius of the PIEZO1 and S2472E structures

The calculation of the curvature radius R of the structure was based on its symmetry and geometric relationship. The distance between the central point of three inner helices and the outermost helix of blade was measured by Chimera as r (radius of the open mouth plane of the bowl). Alternatively, the r was calculated using the equation r = d/√3 by measuring the inter-blade distance between the two outermost helices of two blades. Consistent r value was obtained by using the two different methods. The distance between the plane formed by the outermost helices of three blades and the central site of three inner helices was measured directly by Chimera as h (height of the bowl). Then, the R can be calculated using the equation R^2^=r^2^+(R-h)^2^. All the PIEZO1 and S2472E structures contain 26 out of the 38 TMs. The curvature radius R of PIEZO1 in native membrane at rest was calculated by using the measured inter-blade distance at the distal end of the blade (residue 103) (d = 25 nm)^36^, which gives rise to a r = 25/√3 = 14 nm and R = 23 nm with a presumed h of 5 nm.

### HOLE and CHAP analyses

The central solvent-accessible pathway is generated by the program HOLE. The volumes of CV, EV, MV and IV were calculated using the HOLE data with the formula:

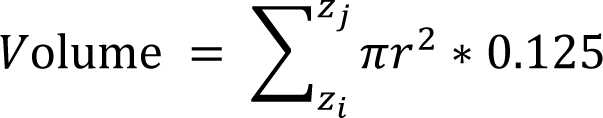

Where 𝑧_𝑖_ and 𝑧_𝑗_ are the lower and upper height limits of the cavity in the z-axis, respectively, *r* is the radius of the hole calculation, and 0.125 Å is the cutoff distance of the hole data in the z-axis. CHAP^21^ analysis was performed according to online instructions (https://www.channotation.org/) and personal communication with Dr. Shanlin Rao. The PDB files of four structures were used as input for CHAP analysis. The output files of this analysis include the time-averaged hydrophobicity due to pore-facing residues and pore radii.

### Molecular dynamics (MD) simulations

MD simulations were performed using the GROMACS2019.6 package^52^ for four PIEZO1 structures: PIEZO1-Curved (PDB: 7WLT) and PIEZO1-Flattened(PDB: 7WLU)^6^, respectively; S2472E-Curved and S2472E-Intermediate were obtained from cryo-EM. The Protein Preparation Wizard module of the Schrödinger2017-2 package was used to predict missing side chains and loops of less than 20 residues. To reduce the size of the simulated system, the structure of PIEZO1 was truncated and only three fragments were retained: residues 1315-1364/1365, 1401/1402/1405-1418/1421 and 1956-2547. This truncated structure was further energy minimized using the AMBER99SB-ILDN^53^ force field with 2000 steps of steepest descent and 3000 steps of a conjugate gradient.

### Coarse-grained system preparation

The truncated structure was converted into a coarse-grained model under the MARTINI v2.2^54^ force field using the martinize.py script, and an elastic network model with a cutoff distance of 0.9 nm was used. The POPC lipid bilayer and solvent were added using the insane.py script, and the system was neutralized using 0.15 M NaCl. The system was then energy minimized by 5000 steps of steepest descent, followed by 5 ns of NVT and 5 ns of NPT simulation to pre-equilibrate, with a time step of 10 fs and POPC beads restrained by a force constant of 1000 kJ • mol^-1^ • nm^-2^. For further pre-equilibration, five successive 5 ns NPT simulations were run with a time step of 20 fs and POPC restrained force constants of 500, 200, 100, 50 and 0 kJ • mol^-1^ • nm^-2^ successive respectively. During NVT and NPT simulations, the protein structure was restrained by a force constant of 1000 kJ • mol^-1^ • nm^-2^. The v-rescale^55^ thermostat was used to couple the protein, lipid, and solvent (water and ions) to a temperature of 310.15 K, while the Berendsen^56^ barostat coupled the pressure to 1 bar of semiisotropic pressure. The cutoff distance of short-range electrostatic and van der Waals interactions was 1.1 nm, while long-range electrostatic interactions were calculated using the reaction-field algorithm. The neighbor list was updated every 20 steps.

### Equilibration and production simulation

The 0.5 µs equilibration simulation was performed to relax the membrane structure, and the positional restraint of the protein was maintained throughout the simulation with a force constant of 1000 kJ • mol^-1^ • nm^-2^. The parameters were identical to those of the NPT equilibration, apart from the change to a Parrinello-Rahman^57^ barostat. At the end of the equilibration simulation, all POPCs that had entered the PIEZO1 pore were removed, and any ions that were present within the protein were exchanged with water outside of the protein, so as to ensure that there were no ions within the protein when the production simulation began. The four PIEZO1 structures were then run for 1 μs at four different transmembrane potentials of 0 V, -0.1 V, -0.25 V, and -0.5 V respectively, and the structures were saved every 0.5 ns. In addition, a 10 μs simulation at 0.5 V of S2472E-Intermediate was performed. The S2472E-Intermediate’s system with the lateral plug domain removed was constructed based on the S2472E-Intermediate’s equilibrated system and also run for 10 μs. During all production simulations, the NC3 and PO4 beads of the POPC and the BB beads of the protein were restrained with a force constant of 500 kJ • mol^-1^ • nm^-2^, which would not allow any change in the protein backbone structure and the lipids could not enter the pore. The transmembrane potential was calculated using 𝑉 = −𝐸𝐿_𝑧_^58^, where 𝐿_𝑧_is the length of the simulated system in the z-axis, *V* is the transmembrane potential difference, and *E* is the electric intensity. The number of Na^+^ in each cavity was counted using MDAnalysis^59^, and the same ion appearing more than 10 times during the simulation was defined as a valid ion. The current of PIEZO1 in the simulation was calculated by the formula:

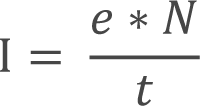

where *e* is the elementary charge, *N* is the number of Na^+^ crossing the PIEZO1 transmembrane channel into the IV region, and *t* is the simulation time. The currents at different potentials were fitted to obtain the slope single-channel conductance. Intracellular conductance was calculated using the number of Na^+^ entering the intracellular space from the IV region. Simulation results were visualized using VMD^60^ (http://www.ks.uiuc.edu/Research/vmd/) and Pymol^61^ (https://github.com/schrodinger/pymol-open-source).

### Statistics and Reproducibility

S2472E proteins were purified more than five times using the method described above. Extended Data Fig. 2b and 4b were representative micrographs of three independent data collections with similar results. Data in Fig. 1, Extended Data Fig. 1 and Extended Data Fig. 5 are shown as mean ± S.E.M. Data normality was tested using the D’Agostino-Pearson omnibus normality test or the Shapiro-Wilk normality test. For those data that follow normal distribution, Statistical significance was evaluated using either unpaired Student’s t-test or ANOVA. Statistical significance was represented as: *p < 0.05, **p < 0.01, ***p < 0.01, ****p < 0.0001.

## Data Availability

The structural coordinates of the Curved and Intermediate PIEZO1 structures were deposited in the Protein Data Bank under the ID 8IXN and 8IXO, respectively. The corresponding cryo-EM maps were deposited into the Electron Microscopy Data Bank (EMDB) under the accession ID EMD-35799 and EMD-35800, respectively.

## Extended Data Figures and Legends

**Extended Data Fig. 1.**
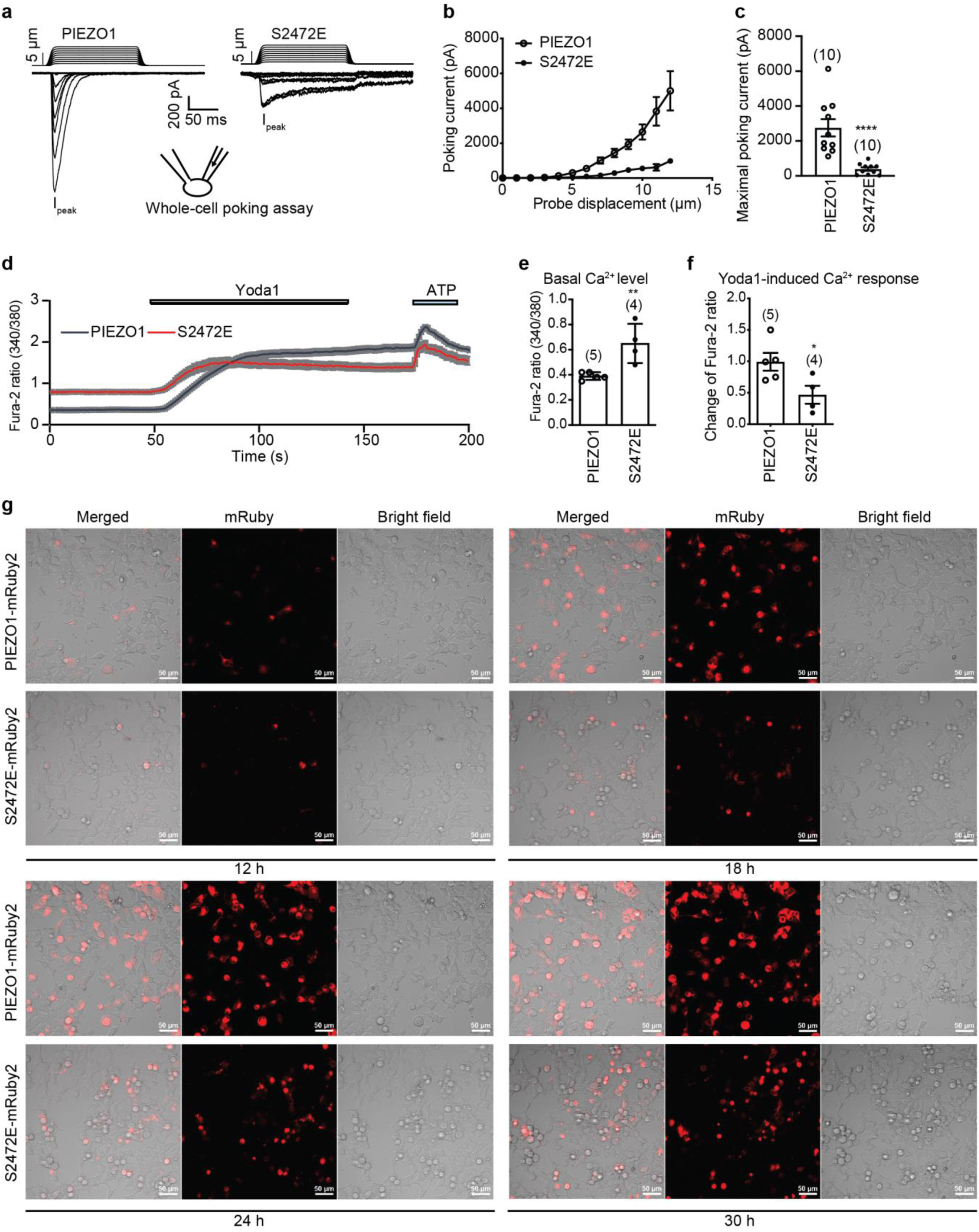
Characterizations of the S2472E mutant. **a**, Representative poking-evoked whole-cell currents of PIEZO1-KO-HEK293T cells transfected with either wild-type mouse PIEZO1 or the S2472E mutant. **b**, Poking induced whole-cell current and probe displacement relationship of PIEZO1-KO-HEK293T cells transfected with either wild-type mouse PIEZO1 or the S2472E mutant. **c**, Scatter plot of the current of the last poking step-evoked whole-cell currents. The recorded cell numbers are labeled. Unpaired Student’s t-test. **P < 0.0001. **d**, Fura-2 calcium imaging traces of PIEZO1 or S2472E-expressing cells show the basal Ca^2+^ level and Yoda1 (5 μM)- or ATP (200 μM)-induced responses. **e**, The basal Ca^2+^ level indicated by the Fura-2 ratio of 340/380 measured from single-cell Ca^2+^ imaging of HEK293T cells transfected with either PIEZO1 or the S2472E mutant. The number of imaged coverslips were labeled. Unpaired Student’s t-test. ****P < 0.01. **f**, The Yoda1-induced change of Fura-2 ratio measured from single-cell Ca^2+^ imaging of HEK293T cells transfected with either PIEZO1 or the S2472E mutant. The number of imaged coverslips were labeled. Unpaired Student’s t-test. *P < 0.05. **g**, Fluorescent and bright field images of HEK293T cell transfected with either PIEZO1-mRuby2 or the S2472E-mRuby2 mutant at the indicated time points. The cells expressing S2472E-mRuby2 became roundup, detached and unhealthy.

**Extended Data Fig. 2.**
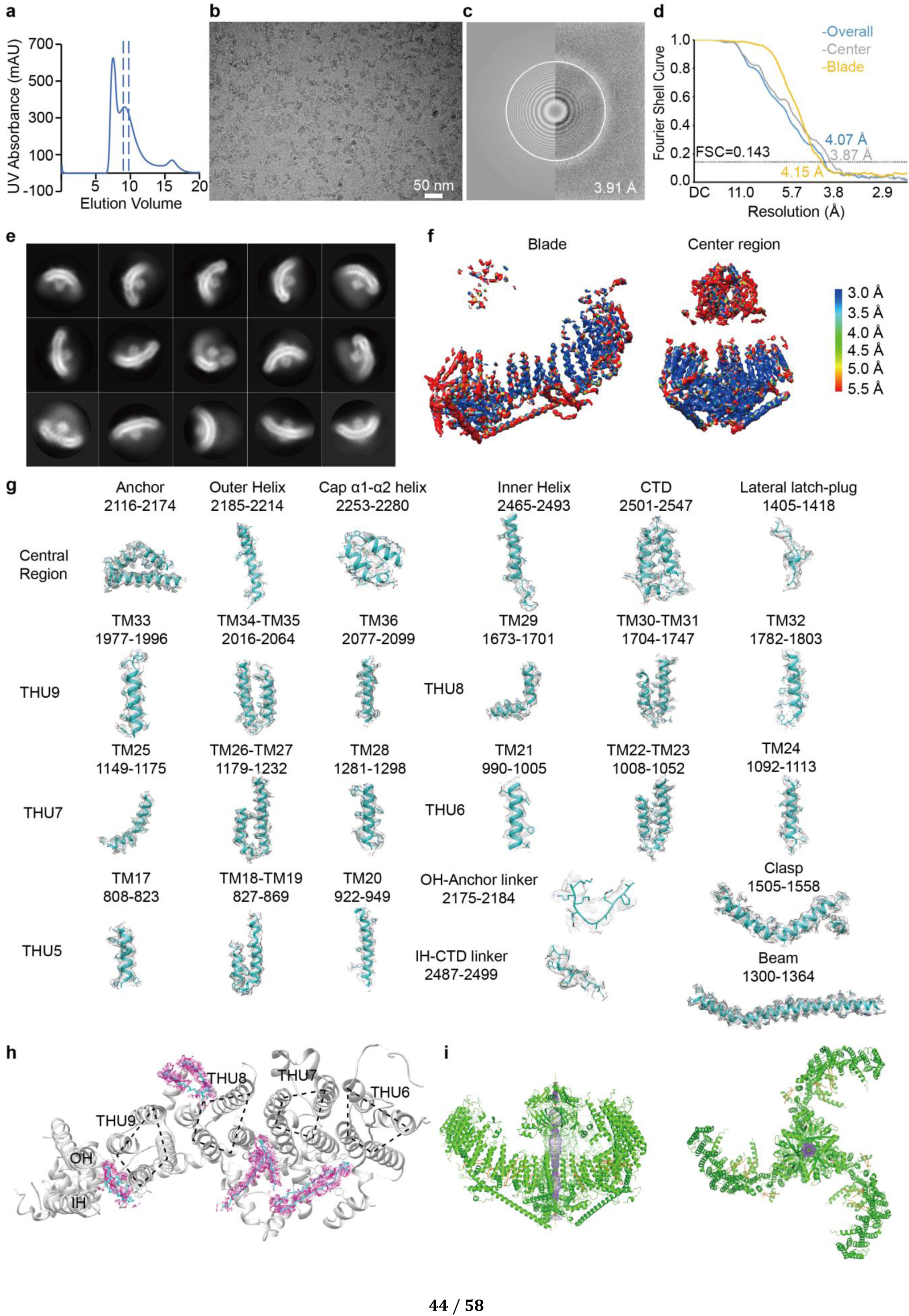
| Cryo-EM determination of the S2472E-Curved structure. **a**, A representative trace of gel filtration of purified S2472E proteins. UV, ultraviolet. Dash lines indicate the peak fractions collected for cryo-sample preparation. **b**, A representative cryo-electron micrograph of mPIEZO1-S2472E solubilized in the detergent GDN. **c**, Power spectrum of the micrograph in **b**, with the 3.91 Å frequency indicated. **d**, Gold-standard Fourier shell correlation (FSC) curves of center, peripheral blade and overall density maps of curved S2472E. Reported resolutions were based on the FSC = 0.143 criterium. **e**, Representative 2D class averages of curved S2472E. **f**, Local-resolution subtracted maps of the S2472E-Curved structure. **g**, Local EM density of the indicated domains of the S2472E-Curved structure. The helices are shown as cartoons with side chains as sticks, fitting with the cryo-EM density shown as gray mesh. **h**, A top view of a single blade of the S2472E-Curved structure, the lipid densities flanking the blade were highlighted in magenta. **i**, The side (left) and top (right) views of the S2472E-Curved structure cartoon model. The pore region was highlighted as purple mesh.

**Extended Data Fig. 3.**
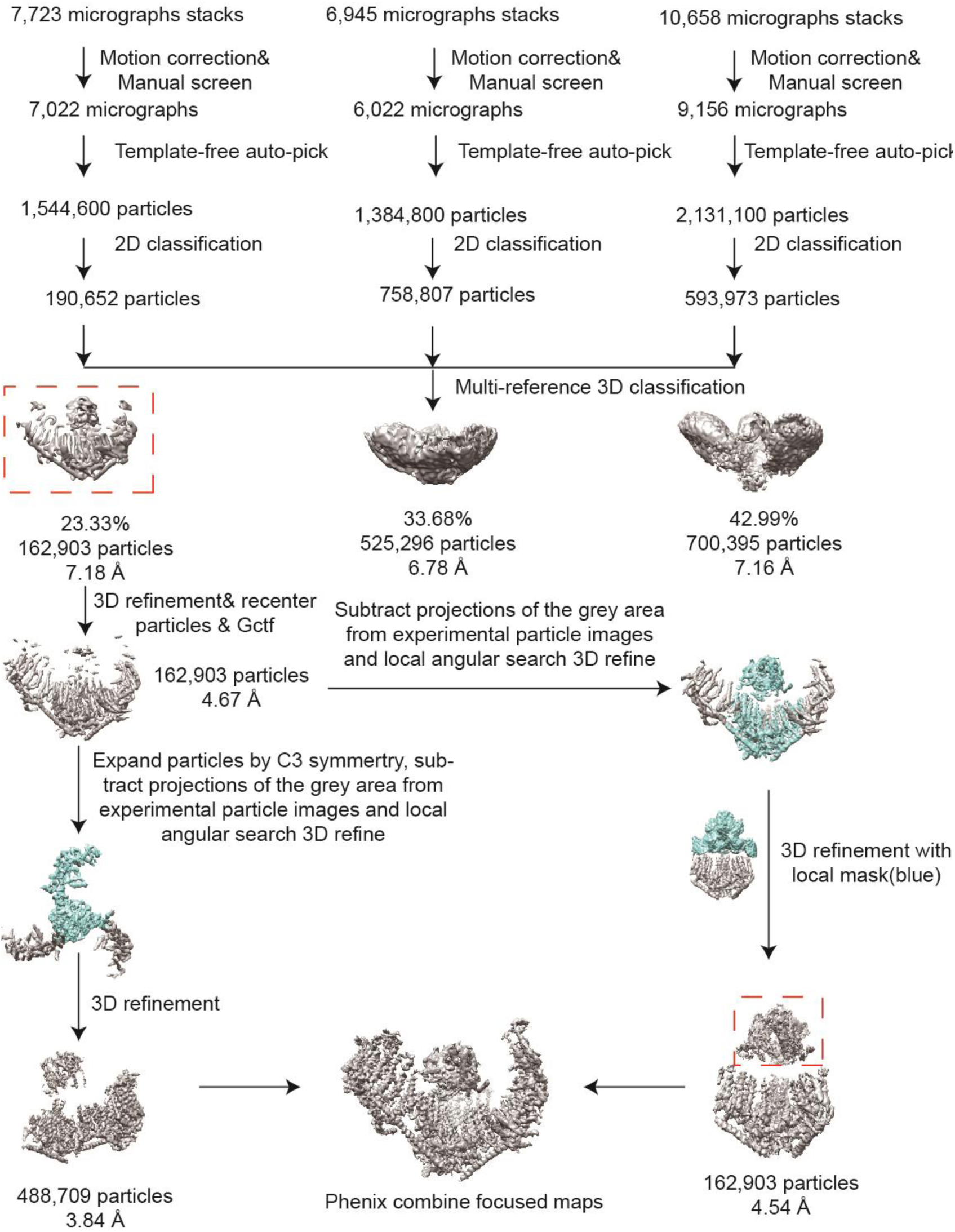
| Flowchart of electron-microscopy data processing of the S2472E-Curved structure. Details of data processing are described in the ‘Imaging processing’ section of the Methods.

**Extended Data Fig. 4.**
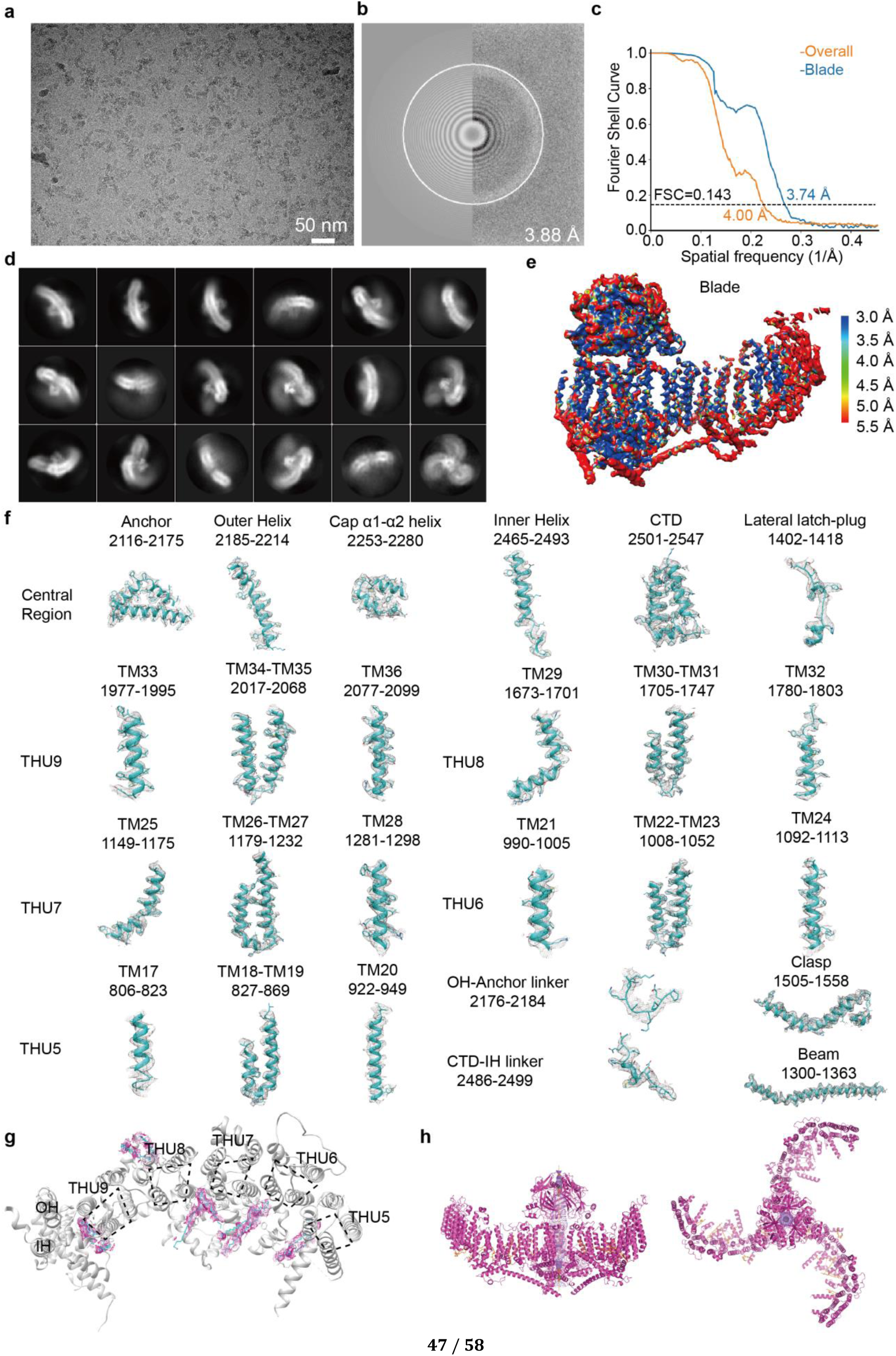
| Cryo-EM determination of the S2472E-Intermediate structure. **a**, A representative cryo-electron micrograph of mPIEZO1-S2472E solubilized in the detergent GDN. **b**, Power spectrum of the micrograph in **a**, with the 3.88 Å frequency indicated. **c**, Gold-standard Fourier shell correlation (FSC) curves of S2472E-Intermediate density map. Reported resolutions were based on the FSC = 0.143 criterium. **d**, Representative 2D class averages of S2472E-Intermediate. **e**, Local-resolution subtracted maps of the S2472E-Intermediate structure. **f**, Local EM density of the indicated domains of the S2472E-Intermediate structure. The helices are shown as cartoons with side chains as sticks, fitting with the cryo-EM density shown as gray mesh. **g**, A top view of a single blade of S2472E-Intermediate structure, the lipid densities flanking the blade were highlighted in magenta. **h**, The side (left) and top (right) views of the S2472E-Intermediate cartoon model. The pore region was highlighted as purple mesh.

**Extended Data Fig. 5.**
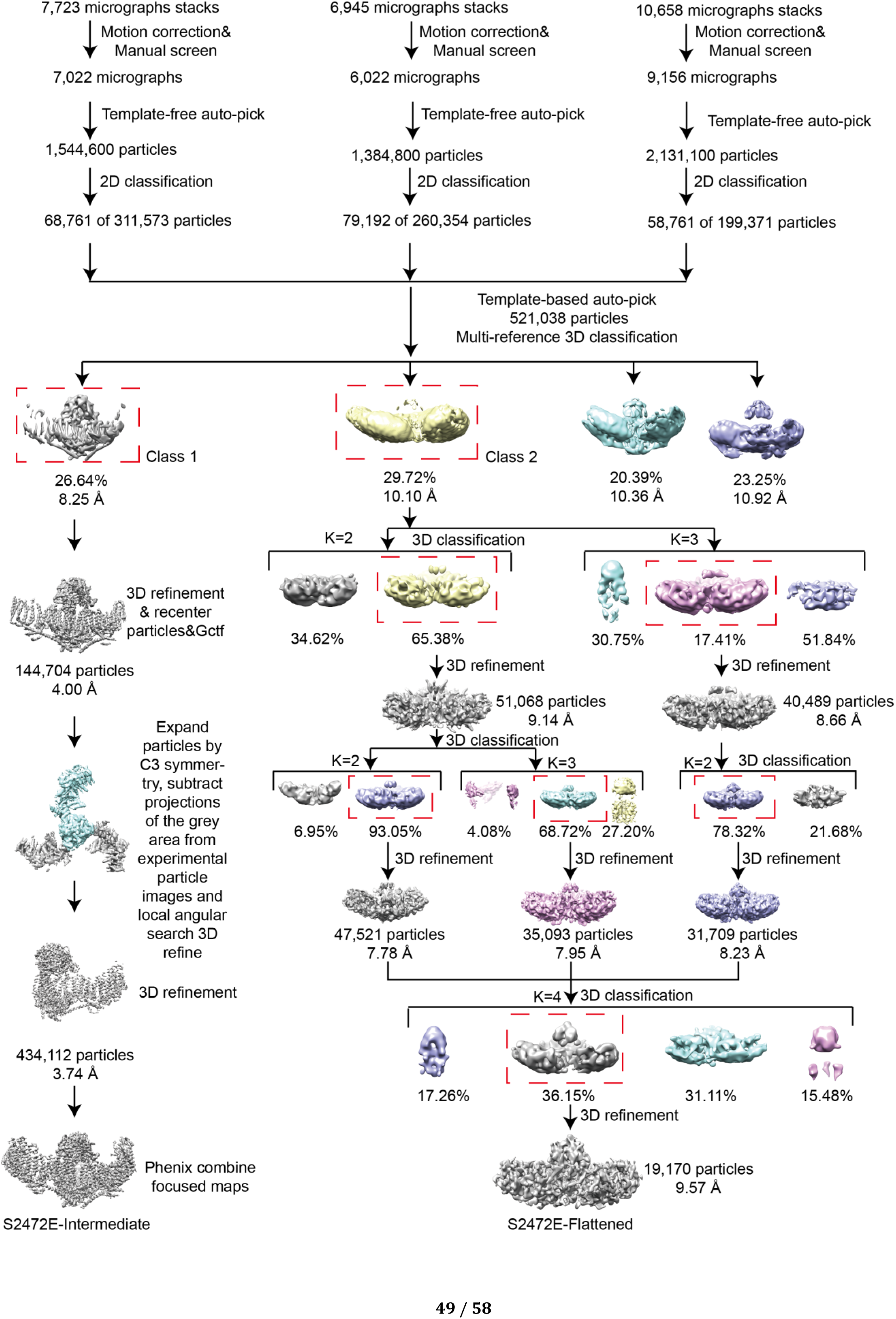
| Flowchart of electron-microscopy data processing of the S2472E-Intermediate and S2472E-Flattened structures. Details of data processing are described in the ‘Imaging processing’ section of the Methods.

**Extended Data Fig. 6.**
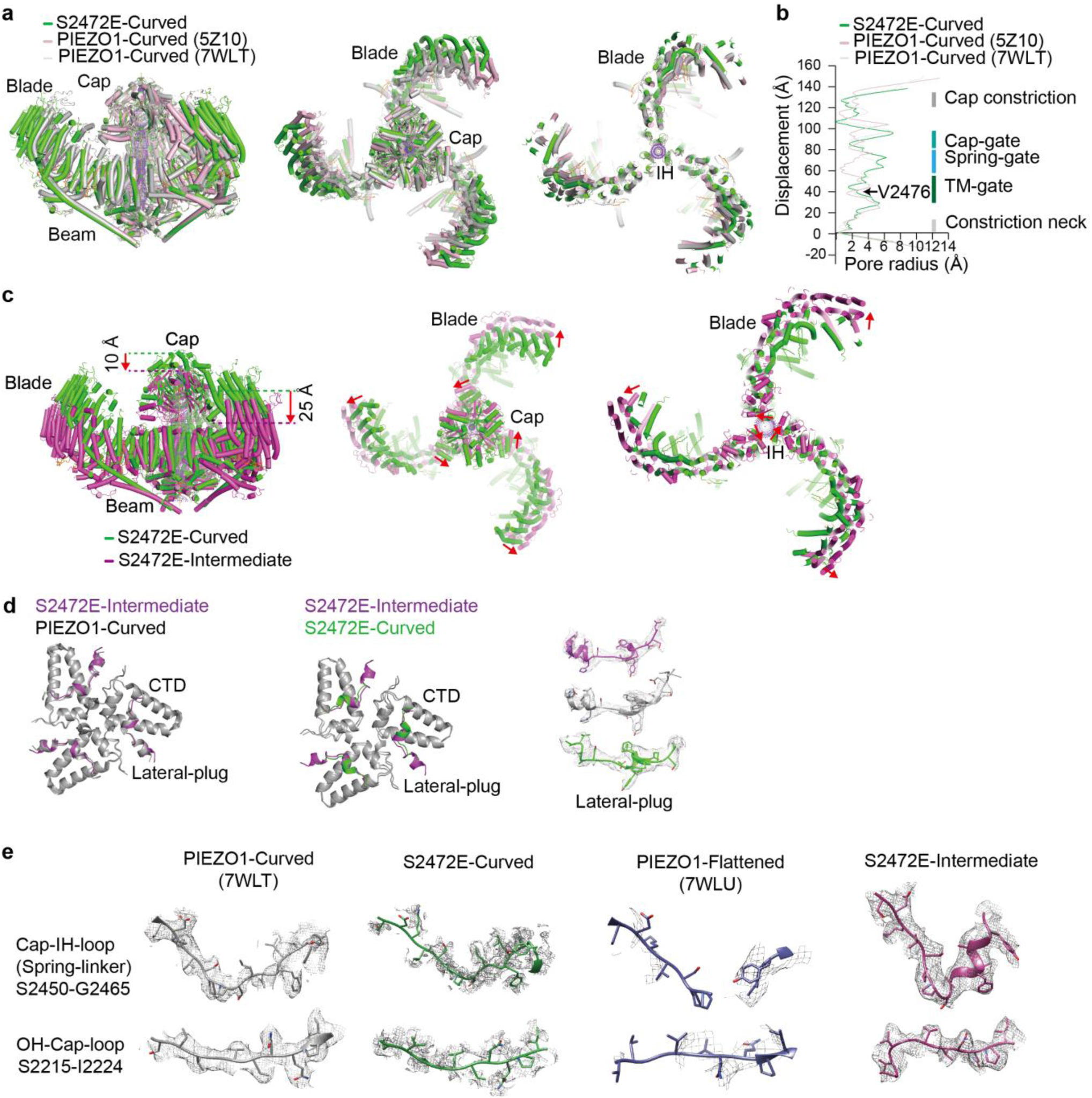
| Structural comparisons. **a**, The overlaid cartoon models of S2472E-Curved structure and PIEZO1-Curved structures derived from either detergent micelles (light pink, PDB: 5Z10) or liposomes (light gray, PDB: 7WLT). The superimposed models were viewed in side (left), or top (middle), or top with the extracellular cap being omitted (right) to clearly display the pore transmembrane helices (IH) and blade transmembrane helices. The Cap, Blade and IH domains are labeled. **b**, Pore radius along the central axis of the ion conduction pathway of the indicated PIEZO1 structures. The positions of Cap constriction, Cap-gate, Spring-gate, TM-gate, and constriction neck are pointed out, and the critical hydrophobic residue V2476 is labeled. **c**, The overlaid cartoon models of S2472E-Curved and S2472E-Intermediate structures. The superimposed models were viewed in side (left), or top (middle), or top with the extracellular cap being omitted (right) to clearly display the pore IH and blade transmembrane helices. The downward motion of Cap and Blade are shown in side view, and the rotation of Cap, Blade, IH are displayed in top view. **d**, The superimposed cartoon models of the central intracellular region including the CTD and the lateral plug of the indicated structures. The lateral plug local EM densities of the S2472E-Intermediate (magenta), PIEZO1-Cureved (gray), and S2472E-curved (green) are displayed for comparison. **e**, The local EM densities of the Cap-IH-loop (spring-linker, upper) or the OH-Cap-loop (lower) of indicated structures.

**Extended Data Fig. 7.**
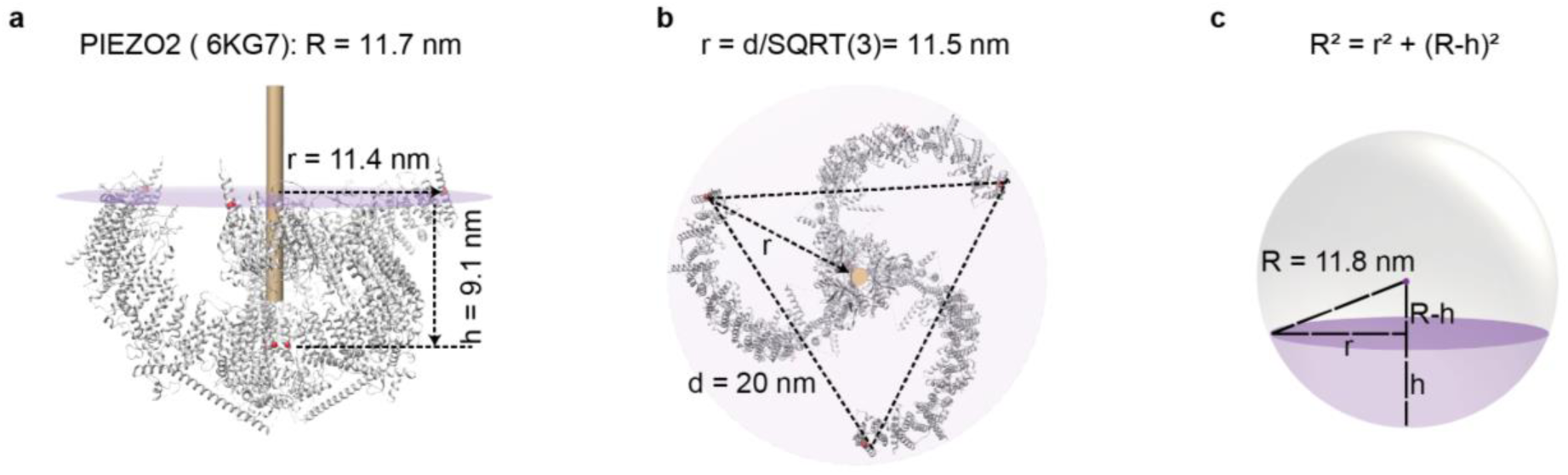
| Measurement of the curvature radius of PIEZO structures using PIEZO2 as an example. **a**, The distance between the central point of three inner helices (indicated by the red dots) and the outermost helix of blade (indicated by the red dots) was measured by Chimera as r (radius of the open mouth plane of the bowl). The distance between the plane formed by the outermost helices of three blades (indicated by the purple plane) and the central site of three inner helices was measured directly by Chimera as h (height of the bowl). **b**, The r was alternatively calculated using the equation r = d/√3 by measuring the inter-blade distance between the two outermost helices of two blades. Consistent r value was obtained by using the two different methods. **c**, The curvature radius R can be calculated using the equation R^2^=r^2^+(R-h)^2^ according to the illustrated relationship among r, h, and R.

**Extended Data Fig. 8.**
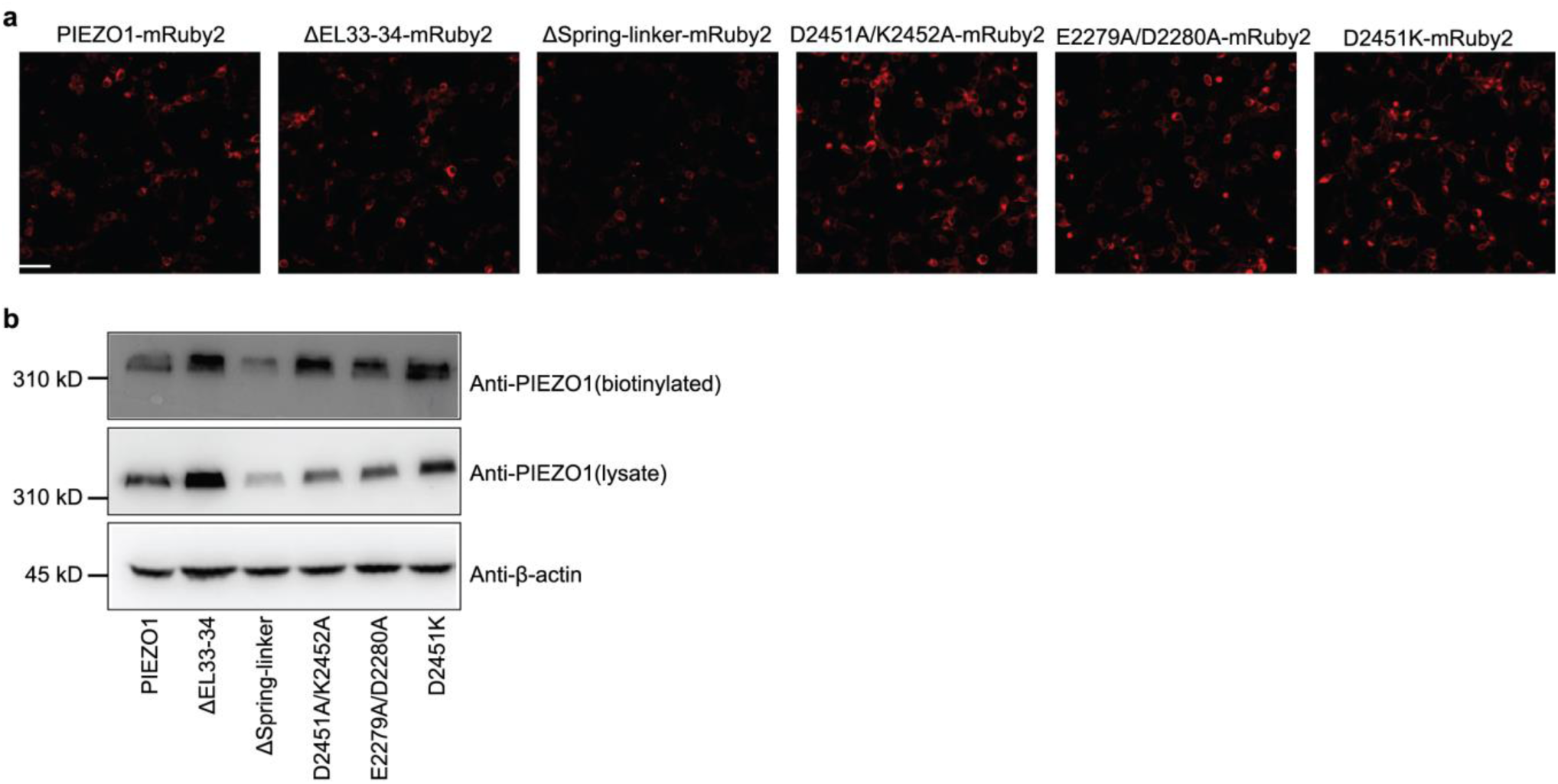
| Expression of PIEZO1 and mutants. **a**, Fluorescent images of HEK293T cells transfected with the indicated PIEZO1 and mutants. **b**, Western blotting of the biotinylated samples (upper) and cell lysate (middle) derived from HEK293T cells transfected with the indicated constructs using the anti-PIEZO1 antibody. The anti-β-actin antibody detected β-actin for a loading control (lower).

**Extended Data Fig. 9.**
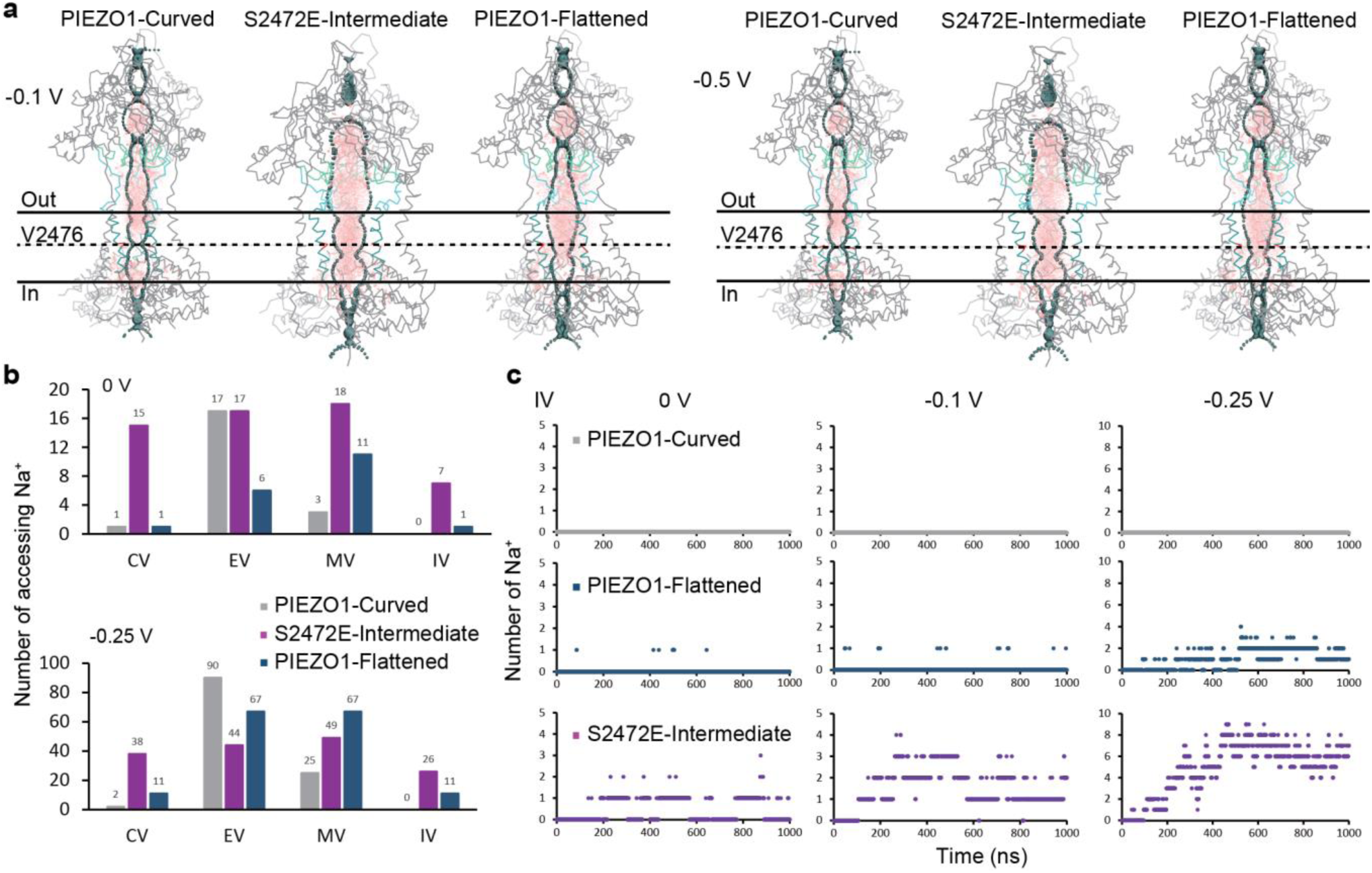
| Molecular dynamic simulations of ion and water permeation. **a**, Molecular dynamic simulations of the water assessing to the CV, EV, MV and IV of the PIEZO1-Curved, S2472E-Intermediate and PIEZO1-Flattened structures. The central pore boundary is illustrated with light teal dots, and the accessing water is represented by the pink dots. The location of the hydrophobic gate residue V2476 is indicated by the black dash line. **b**, The accumulated number of Na^+^ ions accessing the CV, EV, MV and IV of the PIEZO1-Curved (gray), S2472E-Intermediate (purple) and PIEZO1-Flattened (dark blue) at 0 V or -0.25 V. **c**, Time-dependent permeation of Na^+^ to the IV of the PIEZO1-Curved (gray), S2472E-Intermediate (purple) and PIEZO1-Flattened (dark blue) at 0 V, -0.1 V, -0.25.

**Extended Data Table 1.** | Cryo-EM data collection, refinement and validation statistics Cryo-EM data collection, refinement and validation statistics.

|  | #1 Curved state<br>(EMD-35799)<br>(PDB 8IXN) | #2 Intermediate state<br>(EMD-35800)<br>(PDB 8IXO) |
| --- | --- | --- |
| **Data collection and processing** |  |  |
| Magnification | 64,000 | 64,000 |
| Voltage (kV) | 300 | 300 |
| Electron exposure(e-/Å²) | 50 | 50 |
| Defocus range (µm) | -1.5~-2.0 | -1.5~-2.0 |
| Pixel size (Å) | 1.098 | 1.098 |
| Symmetry imposed | C3 | C3 |
| Initial particle images<br>(no.) | 1,543,432 | 521,038 |
| Final particle images<br>(no.) | 488,709 | 434,112 |
| Map resolution (Å) | 4.07 | 4.00 |
| FSC threshold | 0.143 | 0.143 |
| Map resolution range (Å) | 6-3.5 | 10-4.0 |
| **Refinement** |  |  |
| Initial model used (PDB<br>code) | 5Z10 | 5Z10 |
| Model resolution (Å) | 4.3 | 3.9 |
| FSC threshold | 0.5 | 0.5 |
| Map sharpening *B* factor<br>(Å²) | -75 | -68 |
| Model composition |  |  |
| Non-hydrogen atoms | 28521 | 32307 |
| Protein residues | 3966 | 4185 |
| Ligands | 12 | 15 |
| *B* factors (Å²) |  |  |
| Protein | 167.38 | 88.04 |
| Ligand | 105.84 | 111.63 |
| R.m.s. deviations |  |  |
| Bond lengths (Å) | 0.005 | 0.007 |
| Bond angles (°) | 0.776 | 1.120 |
| Validation |  |  |
| MolProbity score | 2.13 | 2.28 |
| Clashscore | 17.09 | 22.01 |
| Poor rotamers (%) | 0.00 | 0.22 |
| Ramachandran plot |  |  |
| Favored (%) | 94.09 | 93.12 |
| Allowed (%) | 5.91 | 6.66 |
| Disallowed (%) | 0.00 | 0.00 |

## Supplementary information

**Supplementary Video 1| Side view of the structural rearrangement from the curved-closed to the intermediate-open state**

**Supplementary Video 2| Top view of the structural rearrangement from the curved-closed to the intermediate-open state**

**Supplementary Video 3 | Side view of the structural rearrangement from the curved-closed to the intermediate-open state of the central pore region**

**Supplementary Video 4 | Top view of the structural rearrangement from the curved-closed to the intermediate-open state of the central pore region**

**Supplementary Video 5| MD simulations of Na^+^ permeation in the PIEZO1-Curved structure**

**Supplementary Video 6| MD simulations of Na^+^ permeation in the S2472E-Intermediate structure**

**Supplementary Video 7| MD simulations of Na^+^ permeation in the PIEZO1-Flattened structure**

### Acknowledgments

We thank Dr. Jie Geng for originally characterizing the S2472E mutant; Drs. Ardem Patapoutian and Eric Mulhall for discussion; the Beijing Advanced Innovation Center for Structural Biology for facility and financial support; and the Protein Preparation and Identification Facility at Technology Center for Protein Science in Tsinghua University for facility support. This work was supported by grant numbers 31825014, 2021ZD0203301, 32130049, 32021002, 31630090, 2016YFA0500402, and 2015CB910102 to B.X.; 31570730, 2016YFA0501102 and 2016YFA0501902 to X.L.; from either the National Natural Science Foundation of China or the National Key R&D Program of China. Also supported by the New Cornerstone Science Foundation through the XPLORER PRIZE, the Research Fund of Vanke School of Public Health and Tsinghua University Initiative Scientific Research Program to B.X., and the Tsinghua-Peking University Center for Life Sciences (20111770319) and the Tsinghua University Initiative Scientific Research Program (20221080048, 20231080030) to B.T.

## Author Contributions

S.L. performed EM sample preparation, data collection, image processing, model building, data analysis and figure preparation; X.Y. carried out protein purification, cryo-EM sample preparation, data collection, cell viability assay and Western blotting; X.C. did molecular cloning, protein purification, cryo-EM sample preparation, Western blotting, immunofluorescent imaging and data collection; X.Z. did MD simulations, data analysis and figure preparation; J.J. did electrophysiology, calcium imaging and data analysis; L.W. helped cryo-EM data collection; J.Y. helped electrophysiological studies; W.L. helped molecular cloning and electrophysiological studies; H.Z. helped model building; K.W. helped electrophysiological studies; B.T. supervised MD simulations studies; X.L. supervised cryo-EM data collection and image processing; B.X. conceived and directed the study, analyzed the structure, made figures and wrote the manuscript with help from all the authors.

## Author Information

The authors declare no competing financial interests. Correspondence and requests for materials should be addressed to B.X., X.L. and B.T.

## References

1 Coste, B. et al. Piezo1 and Piezo2 are essential components of distinct mechanically activated cation channels. Science 330, 55–60, doi:10.1126/science.1193270 (2010).

2 Coste, B. et al. Piezo proteins are pore-forming subunits of mechanically activated channels. Nature 483, 176–181, doi:10.1038/nature10812 (2012).

3 Douguet, D. & Honore, E. Mammalian Mechanoelectrical Transduction: Structure and Function of Force-Gated Ion Channels. Cell 179, 340–354, doi:10.1016/j.cell.2019.08.049 (2019).

4 Kefauver, J. M., Ward, A. B. & Patapoutian, A. Discoveries in structure and physiology of mechanically activated ion channels. Nature 587, 567–576, doi:10.1038/s41586-020-2933-1 (2020).

5 Lin, Y. C. et al. Force-induced conformational changes in PIEZO1. Nature 573, 230–234, doi:10.1038/s41586-019-1499-2 (2019).

6 Yang, X. et al. Structure deformation and curvature sensing of PIEZO1 in lipid membranes. Nature 604, 377–383, doi:10.1038/s41586-022-04574-8 (2022).

7 Syeda, R. et al. Piezo1 Channels Are Inherently Mechanosensitive. Cell Rep 17, 1739–1746, doi:10.1016/j.celrep.2016.10.033 (2016).

8 Cox, C. D. et al. Removal of the mechanoprotective influence of the cytoskeleton reveals PIEZO1 is gated by bilayer tension. Nat Commun 7, 10366, doi:10.1038/ncomms10366 (2016).

9 Xiao, B. Levering Mechanically Activated Piezo Channels for Potential Pharmacological Intervention. Annu Rev Pharmacol Toxicol 60, 195–218, doi:10.1146/annurev-pharmtox-010919-023703 (2020).

10 Del Marmol, J. I., Touhara, K. K., Croft, G. & MacKinnon, R. Piezo1 forms a slowly-inactivating mechanosensory channel in mouse embryonic stem cells. Elife 7, doi:10.7554/eLife.33149 (2018).

11 Peyronnet, R. et al. Piezo1-dependent stretch-activated channels are inhibited by Polycystin-2 in renal tubular epithelial cells. EMBO Rep 14, 1143–1148, doi:10.1038/embor.2013.170 (2013).

12 Shi, J. et al. Sphingomyelinase Disables Inactivation in Endogenous PIEZO1 Channels. Cell Rep 33, 108225, doi:10.1016/j.celrep.2020.108225 (2020).

13 Bae, C., Gnanasambandam, R., Nicolai, C., Sachs, F. & Gottlieb, P. A. Xerocytosis is caused by mutations that alter the kinetics of the mechanosensitive channel PIEZO1. Proc Natl Acad Sci U S A 110, E1162–1168, doi:10.1073/pnas.1219777110 (2013).

14 Albuisson, J. et al. Dehydrated hereditary stomatocytosis linked to gain-of-function mutations in mechanically activated PIEZO1 ion channels. Nat Commun 4, 1884, doi:10.1038/ncomms2899 (2013).

15 Ge, J. et al. Architecture of the mammalian mechanosensitive Piezo1 channel. Nature 527, 64–69, doi:10.1038/nature15247 (2015).

16 Wang, L. et al. Structure and mechanogating of the mammalian tactile channel PIEZO2. Nature 573, 225–229, doi:10.1038/s41586-019-1505-8 (2019).

17 Zhao, Q. et al. Structure and mechanogating mechanism of the Piezo1 channel. Nature 554, 487–492, doi:10.1038/nature25743 (2018).

18 Saotome, K. et al. Structure of the mechanically activated ion channel Piezo1. Nature 554, 481–486, doi:10.1038/nature25453 (2018).

19 Guo, Y. R. & MacKinnon, R. Structure-based membrane dome mechanism for Piezo mechanosensitivity. Elife 6, doi:10.7554/eLife.33660 (2017).

20 Rao, S., Klesse, G., Stansfeld, P. J., Tucker, S. J. & Sansom, M. S. P. A heuristic derived from analysis of the ion channel structural proteome permits the rapid identification of hydrophobic gates. Proc Natl Acad Sci U S A 116, 13989–13995, doi:10.1073/pnas.1902702116 (2019).

21 Klesse, G., Rao, S., Sansom, M. S. P. & Tucker, S. J. CHAP: A Versatile Tool for the Structural and Functional Annotation of Ion Channel Pores. Journal of Molecular Biology 431, 3353–3365, 10.1016/j.jmb.2019.06.003 (2019).

22 Zheng, W., Gracheva, E. O. & Bagriantsev, S. N. A hydrophobic gate in the inner pore helix is the major determinant of inactivation in mechanosensitive Piezo channels. Elife 8, doi:10.7554/eLife.44003 (2019).

23 Syeda, R. et al. Chemical activation of the mechanotransduction channel Piezo1. Elife 4, doi:10.7554/eLife.07369 (2015).

24 Aryal, P., Sansom, M. S. & Tucker, S. J. Hydrophobic gating in ion channels. J Mol Biol 427, 121–130, doi:10.1016/j.jmb.2014.07.030 (2015).

25 Geng, J. et al. A Plug-and-Latch Mechanism for Gating the Mechanosensitive Piezo Channel. Neuron 106, 438–451 e436, doi:10.1016/j.neuron.2020.02.010 (2020).

26 Jiang, W. et al. Crowding-induced opening of the mechanosensitive Piezo1 channel in silico. Communications Biology 4, 84, doi:10.1038/s42003-020-01600-1 (2021).

27 Wu, J. et al. Inactivation of Mechanically Activated Piezo1 Ion Channels Is Determined by the C-Terminal Extracellular Domain and the Inner Pore Helix. Cell Rep 21, 2357–2366, doi:10.1016/j.celrep.2017.10.120 (2017).

28 Lewis, A. H. & Grandl, J. Inactivation Kinetics and Mechanical Gating of Piezo1 Ion Channels Depend on Subdomains within the Cap. Cell Rep 30, 870–880.e872, doi:10.1016/j.celrep.2019.12.040 (2020).

29 Zhao, Q. et al. Ion Permeation and Mechanotransduction Mechanisms of Mechanosensitive Piezo Channels. Neuron 89, 1248–1263, doi:10.1016/j.neuron.2016.01.046 (2016).

30 Peralta, F. A. et al. Optical control of PIEZO1 channels. Nat Commun 14, 1269, doi:10.1038/s41467-023-36931-0 (2023).

31 Jiang, W., Lacroix, J. & Luo, Y. L. Importance of molecular dynamics equilibrium protocol on protein-lipid interaction near channel pore. Biophys Rep (N Y) 2, 100080, doi:10.1016/j.bpr.2022.100080 (2022).

32 Wang, Y. et al. A lever-like transduction pathway for long-distance chemical- and mechano-gating of the mechanosensitive Piezo1 channel. Nature Communications 9, 1300, doi:10.1038/s41467-018-03570-9 (2018).

33 Lewis, A. H. & Grandl, J. Mechanical sensitivity of Piezo1 ion channels can be tuned by cellular membrane tension. Elife 4, doi:10.7554/eLife.12088 (2015).

34 Haselwandter, C. A., Guo, Y. R., Fu, Z. & MacKinnon, R. Quantitative prediction and measurement of Piezo’s membrane footprint. Proc Natl Acad Sci U S A 119, e2208027119, doi:10.1073/pnas.2208027119 (2022).

35 Haselwandter, C. A., Guo, Y. R., Fu, Z. & MacKinnon, R. Elastic properties and shape of the Piezo dome underlying its mechanosensory function. Proc Natl Acad Sci U S A 119, e2208034119, doi:10.1073/pnas.2208034119 (2022).

36 Mulhall, E. M. et al. Direct observation of the conformational states of PIEZO1. Nature, doi:10.1038/s41586-023-06427-4 (2023).

37 Xiao, B., Coste, B., Mathur, J. & Patapoutian, A. Temperature-dependent STIM1 activation induces Ca2+ influx and modulates gene expression. Nature Chemical Biology 7, 351–358, doi:10.1038/nchembio.558 (2011).

38 Wang, J. et al. Tethering Piezo channels to the actin cytoskeleton for mechanogating via the cadherin-beta-catenin mechanotransduction complex. Cell Rep 38, 110342, doi:10.1016/j.celrep.2022.110342 (2022).

39 Baxa, U. Imaging of Liposomes by Transmission Electron Microscopy. Methods in molecular biology (Clifton, N.J.) 1682, 73–88, doi:10.1007/978-1-4939-7352-1_8 (2018).

40 Zheng, S. Q. et al. MotionCor2: anisotropic correction of beam-induced motion for improved cryo-electron microscopy. Nature methods 14, 331–332, doi:10.1038/nmeth.4193 (2017).

41 Mindell, J. A. & Grigorieff, N. Accurate determination of local defocus and specimen tilt in electron microscopy. Journal of structural biology 142, 334–347, doi:10.1016/s1047-8477(03)00069-8 (2003).

42 Zivanov, J. et al. New tools for automated high-resolution cryo-EM structure determination in RELION-3. eLife 7, doi:10.7554/eLife.42166 (2018).

43 Zhang, K. Gctf: Real-time CTF determination and correction. J Struct Biol 193, 1–12, doi:10.1016/j.jsb.2015.11.003 (2016).

44 Pettersen, E. F. et al. UCSF Chimera--a visualization system for exploratory research and analysis. Journal of computational chemistry 25, 1605–1612, doi:10.1002/jcc.20084 (2004).

45 Adams, P. D. et al. PHENIX: a comprehensive Python-based system for macromolecular structure solution. *Acta crystallographica. Section D*, Biological crystallography 66, 213–221, doi:10.1107/s0907444909052925 (2010).

46 Scheres, S. H. & Chen, S. Prevention of overfitting in cryo-EM structure determination. Nat Methods 9, 853–854, doi:10.1038/nmeth.2115 (2012).

47 Rosenthal, P. B. & Henderson, R. Optimal determination of particle orientation, absolute hand, and contrast loss in single-particle electron cryomicroscopy. Journal of molecular biology 333, 721–745, doi:10.1016/j.jmb.2003.07.013 (2003).

48 Kucukelbir, A., Sigworth, F. J. & Tagare, H. D. Quantifying the local resolution of cryo-EM density maps. Nat Methods 11, 63–65, doi:10.1038/nmeth.2727 (2014).

49 Punjani, A., Rubinstein, J. L., Fleet, D. J. & Brubaker, M. A. J. N. M. cryoSPARC: algorithms for rapid unsupervised cryo-EM structure determination. 14, 290–296 (2017).

50 Emsley, P., Lohkamp, B., Scott, W. G. & Cowtan, K. Features and development of Coot. Acta crystallographica. Section D, Biological crystallography 66, 486–501, doi:10.1107/S0907444910007493 (2010).

51 Williams, C. J. et al. MolProbity: More and better reference data for improved all-atom structure validation. Protein science : a publication of the Protein Society 27, 293–315, doi:10.1002/pro.3330 (2018).

52 Abraham, M. J. et al. GROMACS: High performance molecular simulations through multi-level parallelism from laptops to supercomputers. SoftwareX 1-2, 19–25, 10.1016/j.softx.2015.06.001 (2015).

53 Lindorff-Larsen, K. et al. Improved side-chain torsion potentials for the Amber ff99SB protein force field. Proteins 78, 1950–1958, doi:10.1002/prot.22711 (2010).

54 de Jong, D. H. et al. Improved Parameters for the Martini Coarse-Grained Protein Force Field. Journal of Chemical Theory and Computation 9, 687–697, doi:10.1021/ct300646g (2013).

55 Bussi, G., Donadio, D. & Parrinello, M. Canonical sampling through velocity rescaling. The Journal of chemical physics 126, 014101, doi:10.1063/1.2408420 (2007).

56 Berendsen, H. J. C., Postma, J. P. M., Gunsteren, W. F. v., DiNola, A. & Haak, J. R. Molecular dynamics with coupling to an external bath. The Journal of chemical physics 81, 3684–3690, doi:10.1063/1.448118 (1984).

57 Parrinello, M. & Rahman, A. Polymorphic transitions in single crystals: A new molecular dynamics method. Journal of Applied Physics 52, 7182–7190, doi:10.1063/1.328693 (1981).

58 Aksimentiev, A. & Schulten, K. Imaging α-Hemolysin with Molecular Dynamics: Ionic Conductance, Osmotic Permeability, and the Electrostatic Potential Map. Biophysical Journal 88, 3745–3761, 10.1529/biophysj.104.058727 (2005).

59 Michaud-Agrawal, N., Denning, E. J., Woolf, T. B. & Beckstein, O. MDAnalysis: a toolkit for the analysis of molecular dynamics simulations. J Comput Chem 32, 2319–2327, doi:10.1002/jcc.21787 (2011).

60 Humphrey, W., Dalke, A. & Schulten, K. VMD: Visual molecular dynamics. Journal of Molecular Graphics 14, 33–38, 10.1016/0263-7855(96)00018-5 (1996).

61 DeLano, W. L. Pymol: An open-source molecular graphics tool. CCP4 Newsl. Protein Crystallogr 40, 82–92 (2002).

